# Metabolic regulation of species-specific developmental rates

**DOI:** 10.1101/2021.08.27.457974

**Authors:** Margarete Diaz-Cuadros, Teemu P. Miettinen, Dylan Sheedy, Carlos Manlio Díaz-García, Svetlana Gapon, Alexis Hubaud, Gary Yellen, Scott R. Manalis, William Oldham, Olivier Pourquié

**Affiliations:** Department of Genetics, Harvard Medical School and Department of Pathology, Brigham and Women’s Hospital, Boston, MA, USA; Koch Institute for Integrative Cancer Research, Massachusetts Institute of Technology, Cambridge, MA, USA; Department of Neurobiology, Harvard Medical School, Boston, MA, USA; Department of Biological Engineering, Massachusetts Institute of Technology, Cambridge, MA, USA; Department of Mechanical Engineering, Massachusetts Institute of Technology, Cambridge, MA, USA; Department of Medicine, Harvard Medical School and Department of Medicine, Brigham and Women’s Hospital, Boston, MA, USA; Harvard Stem Cell Institute, Harvard University, Cambridge, MA USA

## Abstract

Animals display significant inter-specific variation in the rate of embryonic development despite broad conservation of the overall sequence of developmental events. Differences in biochemical reaction speeds, including the rates of protein production and degradation, are thought to be responsible for distinct species-specific rates of development [1–3]. However, the cause of differential biochemical reaction speeds between species remains unknown. Using pluripotent stem cells, we have established an *in vitro* system that recapitulates the two-fold difference in developmental rate between early mouse and human embryos. This system provides a quantitative measure of developmental speed as revealed by the period of the segmentation clock, a molecular oscillator associated with the rhythmic production of vertebral precursors. Using this system, we showed that mass-specific metabolic rates scale with developmental rate and are therefore elevated in mouse cells compared to human cells. We further showed that reducing these metabolic rates by pharmacologically inhibiting the electron transport chain slows down the segmentation clock. The effect of the electron transport chain on the segmentation clock is mediated by the cellular NAD^+^/NADH redox balance independent of ATP production and, further downstream, by the global rate of protein synthesis. These findings represent a starting point for the manipulation of developmental rate, which would find multiple translational applications including the acceleration of human pluripotent stem cell differentiation for disease modeling and cell-based therapies.

---

The rate of embryonic development across animal taxa is a highly variable and species-specific trait that is correlated with lifespan, body size, and other life history traits [4, 5]. In mammals, large-bodied species develop at slower rates and display increased lifespans (e.g. humans) compared to small-bodied animals (e.g. mice) [6]. Astonishingly, the duration of gestation in mammals ranges from 20 days in mice to 645 days in African elephants [7, 8]. Differences in developmental rate are best illustrated at early stages of embryogenesis when different species are most morphologically and molecularly similar. At 37°C, mouse embryos start gastrulation at day 7 post-fertilization (PF), the first somite forms at day 8, and the forelimb bud is first visible at day 9; whereas in human embryos gastrulation begins at day 16 PF, the first somite forms at day 20, and the forelimb bud is first seen at day 30 [9]. Thus, even though early mouse and human embryos undergo the same series of developmental steps and share a similar overall size, human embryos do so at rates 2-3 times slower [9].

Such differences in developmental rate between mouse and human embryos can be recapitulated *in vitro* by studying the differentiation kinetics of pluripotent stem cells (PSCs) [1, 2, 10, 11]. We previously reported the establishment of PSC-derived models of the mouse and human segmentation clocks, which displayed approximately two-fold slower oscillations in human cells compared to mouse cells [12]. The segmentation clock is a molecular oscillator that operates in presomitic mesoderm (PSM) cells and controls the periodicity of somite formation in vertebrate embryos [13]. This clock represents an ideal model for developmental rate because its period is species-specific, temperature-sensitive, and scales with the speed of development [1, 14, 15]. We thus adopted our *in vitro* segmentation clock system as a platform to directly compare developmental rate between mouse and human cells.

To mitigate the influence of potential confounding factors in the comparison of mouse and human cells, we first developed an updated protocol that allowed mouse and human PSCs to be differentiated towards PSM fate under identical conditions (Fig. 1a). This protocol still relies on Wnt activation coupled with BMP inhibition to induce PSM, but the basal media was standardized for both species. The differentiation efficiency, as measured by induction of a specific posterior PSM fate marker (MSGN1-Venus), was remarkably high at 78.3 ± 1% for mouse cells and 96.5 ± 1.5% for human cells (Fig. 1b, Extended Data Fig. 1a). Under these conditions, we observed differences in developmental rate between mouse and human cells at several levels. First, mouse cells activated MSGN1-Venus with accelerated kinetics compared to human cells (1-2 days for mouse; 2-3 days for human; Fig. 1b). Second, the cell cycle duration was shorter in mouse (13.9 ± 2 hours) than in human (21.9 ± 3.6 hours) PSM cells (Fig. 1c). Lastly, as previously reported, oscillations of the segmentation clock reporter HES7-Achilles were faster in mouse (2.6 ± 0.3 hours) compared to human (4.9 ± 0.3 hours) PSM cells (Fig. 1d-e) [12, 16–18]. Together, these observations indicated an approximately two-fold difference in developmental rate between mouse and human PSM cells differentiating *in vitro*.

**Figure 1.**
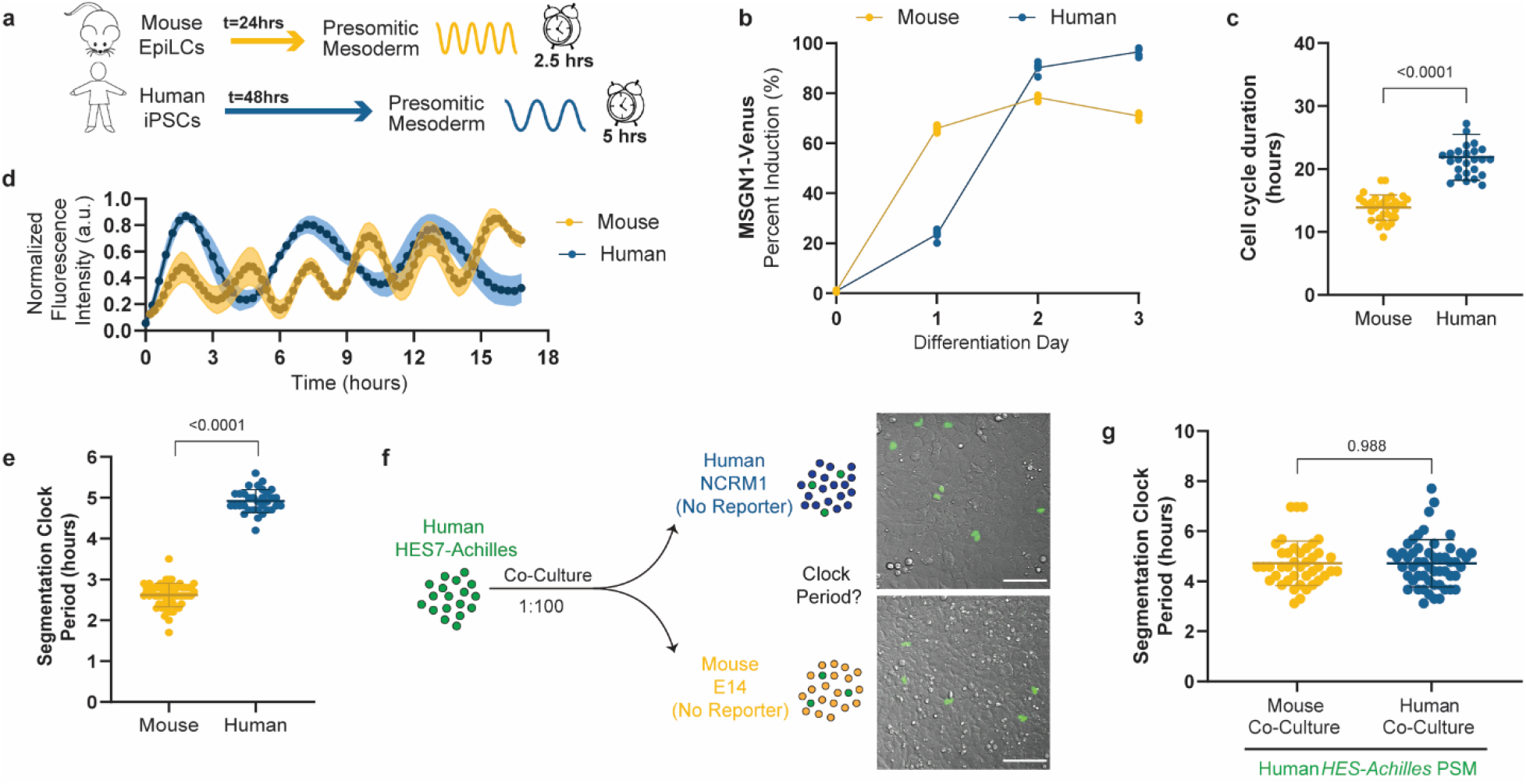
Cell-autonomous differences in developmental rate between differentiating mouse and human presomitic mesoderm (PSM) cells a. Schematic illustrating the differentiation of mouse and human PSCs towards PSM fate. The accelerated developmental pace of mouse cells is reflected in the reduced induction time and short oscillatory period relative to human cells. EpiLCs = Epliblast-Like Cells, iPSCs = induced Pluripotent Stem Cells. b. PSM induction efficiency over the course of 3 days of differentiation for mouse and human PSCs. The percentage of cells expressing MSGN1-Venus was assessed by flow cytometry. n=5. c. Duration of the cell cycle in hours for PSC-derived mouse and human PSM cells. Mean ±SD. n=33 (mouse); n=26 (human). d. HES7-Achilles oscillation profiles for PSC-derived mouse and human PSM cells over the course of 18 hours. Mean ±SEM. n=5. e. Period of HES7-Achilles oscillations in PSC-derived mouse and human PSM cells. Mean ±SD. n=25. f. Left: Experimental strategy for the co-culture of *HES7-Achilles* PSM human cells with non-reporter mouse (*E14*) or human (*NCRM1*) PSM cells at a ratio of 1:100. Right: Merged brightfield and HES7-Achilles fluorescence images of human-human (top) and human-mouse (bottom) co-cultures. Scale bar = 100μm. g. Period of HES7-Achilles oscillations in human PSM cells co-cultured with either mouse or human PSM cells. Mean ±SD. n=41 (mouse co-culture), n=53 (human co-culture).

To validate that this *in vitro* model accurately recapitulated developmental rate, we directly compared the cell cycle time and segmentation clock period of PSC-derived mouse PSM cells with primary explants of mouse PSM taken from E9.5 embryos [19]. We found that these parameters did not significantly differ between PSC-derived and primary PSM cells (Extended Data Fig. 1b-c). These results confirmed that PSM cells generated *in vitro* can precisely reflect the endogenous pace of development.

Differences in developmental rate can be cell autonomous or context dependent. For instance, the neural differentiation rate of human PSCs can be accelerated by co-culturing them with an excess of mouse cells [20]. In contrast, human PSC derivatives maintain their slow developmental rate during teratoma formation in a mouse host or when grafted as cortical neurons into the mouse brain [10, 21]. The segmentation clock retains its species-specific period in isolated cells, thus suggesting that it is a cell autonomous feature that does not require tissue-level coupling [1, 12, 19]. To further determine whether the period of the segmentation clock is indeed cell autonomous, we co-cultured *HES7-Achilles* human PSM cells with control PSM cells of either human or mouse origin at a ratio of 1:100 (Fig. 1f). The segmentation clock period was the same when co-cultured with either human (4.71 ± 0.94 hours) or mouse (4.71 ±0. 88 hours) PSM cells (Fig. 1g). This result indicated that the segmentation clock period is cell autonomous even in chimeric conditions.

The cell cycle has been proposed to function as a clock that controls developmental speed [2, 22]. Our observation that cell cycle duration scales with developmental rate in mouse and human PSM cells motivated us to test this hypothesis. We blocked cell division in human PSM cells with the replication inhibitor aphidicolin. However, complete arrest of the cell cycle at early S phase did not impact the oscillatory period (Fig. Extended Data Fig. 1d-e). We thus conclude that the cell cycle does not contribute to the regulation of the segmentation clock period.

In general, the mass-specific metabolic rate of smaller animals is accelerated compared to that of larger animals [23, 24]. Kleiber’s law, a well-established allometric relationship between body mass and metabolic rate, applies to most animals (Kleiber’s law) [23]. Whether this reflects scaling of the intrinsic cellular metabolic rates or constraints in energy delivery by the vascular system in adults is controversial [24, 25]. The cellular mass-specific metabolic rate has not been compared across species during development and thus whether metabolic rate scales with developmental rate remains unknown. Such differences in basal metabolism could provide an attractive hypothesis to explain the accelerated biochemical kinetics observed in mouse cells compared to human cells [1, 2].

Having established that our *in vitro* segmentation clock system represented a reliable and cell-autonomous model of developmental rate, we set out to compare the rates of metabolism of mouse and human PSM cells directly. We first measured the oxygen consumption rate (OCR) and glycolytic proton efflux rate (glycoPER). When an identical number of cells were seeded per well, no significant difference was observed in basal glycoPER between species, and basal OCR was slightly elevated in human cells (Extended Data Fig. 2a-b). However, we noted that human PSM cells were approximately twice as large in volume as mouse cells (2060 ± 524 fL vs. 885.9 ± 157.9 fL, respectively) (Fig. 2a; Extended Data Fig. 2c). To compare how total cell mass differs between human and mouse cells, we measured the total mass, volume and density of individual cells by repeatedly weighing the cells in different density fluids using the suspended microchannel resonator [26, 27]. The total mass of human cells was also approximately twice that of mouse cells (2002 ± 71 pg vs. 1066 ± 56 pg) (Fig. 2b), such that the overall density was similar for both species (1.057 ± 0.003 pg/fL vs. 1.061 ± 0.002 pg/fL; Fig. 2c). PSC-derived human PSM cells were thus two-fold larger and heavier than mouse PSM cells. Therefore, the metabolic rate per cell could not be directly compared between mouse and human PSM cells. Instead, we compared the mass-specific metabolic rates and found that mass-specific OCR and glycoPER were twice as fast in mouse cells as in human cells (Fig. 2d-e).

**Figure 2.**
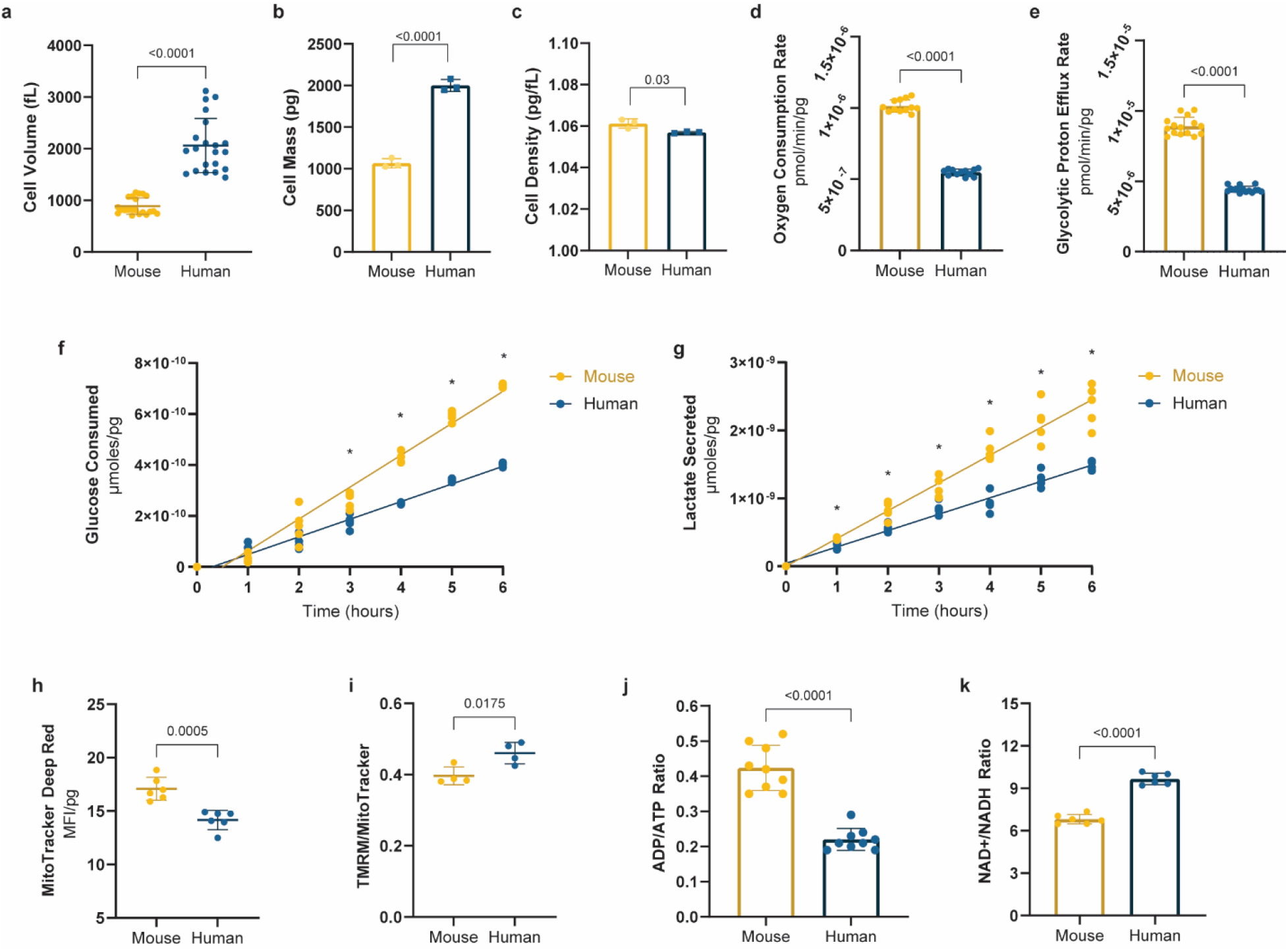
Elevated mass-specific metabolic rates in mouse PSM cells compared to human PSM cells a. Volume of MSGN1-Venus+ PSC-derived mouse and human PSM cells as measured with a coulter counter. Mean ±SD. n=21. b. Total cell mass of MSGN1-Venus+ PSC-derived mouse and human PSM cells as measured on a suspended microchannel resonator. Mean ±SD. n=3. c. Total cell density of MSGN1-Venus+ PSC-derived mouse and human PSM cells as measured on a suspended microchannel resonator. Mean ±SD. n=3. d. Mass-specific oxygen consumption rate for MSGN1-Venus+ PSC-derived mouse and human PSM cells. Mean ±SD. n=12. e. Mass-specific glycolytic proton efflux rate for MSGN1-Venus+ PSC-derived mouse and human PSM cells. Mean ±SD. n=15. f. Mass-specific cumulative glucose consumption over the course of 6 hours for MSGN1-Venus+ PSC-derived mouse and human PSM cells. Each datapoint represents the average of technical duplicates. n=5. g. Mass-specific cumulative lactate secretion over the course of 6 hours for MSGN1-Venus+ PSC-derived mouse and human PSM cells. Each datapoint represents the average of technical duplicates. n=5. h. Mass-specific mitochondrial content in PSC-derived mouse and human MSGN1-Venus+ PSM cells as measured with the MitoTracker Deep Red dye and flow cytometry. Mean ±SD. n=6. i. Inner mitochondrial membrane potential (Δψm) in MSGN1-Venus+ PSC-derived mouse and human PSM cells. TMRM fluorescence was normalized by mitochondrial content following flow cytometry. Mean ±SD. n=4. j. Whole-cell ADP/ATP ratio in MSGN1-Venus+ PSC-derived mouse and human PSM cells. Mean ±SD. n=9. Each datapoint represents the average of 3 technical replicates. k. Whole-cell NAD^+^/NADH ratio in MSGN1-Venus+ PSC-derived mouse and human PSM cells. Each datapoint represents the average of 3 technical replicates. Mean ±SD. n=6.

We confirmed that mass-specific metabolic rates are faster in mouse cells than in human cells using multiple independent approaches. We measured glucose consumption and lactate secretion over the course of six hours for both species. Both glucose consumption and lactate secretion per unit mass were significantly higher in mouse cells compared to human cells (Fig. 2f-g). The mass-specific glutamine consumption, which fuels the TCA cycle through glutaminolysis, was also significantly elevated in mouse PSM cells (Extended Data Fig. 2d). We performed stable isotope tracing with uniformly labeled U-^13^C_6_-Glucose and U-^13^C_5_-Glutamine in both mouse and human PSM cells. At isotopic steady state, we observed that glucose tracing led to high levels of labeling for pyruvate and lactate, as well as partial labeling of TCA intermediates (Extended Data Fig. 2e-j). Glutamine tracing showed intermediate levels of labelling for glutamate and TCA metabolites, but not pyruvate nor lactate supporting an anaplerotic role in the TCA cycle (Extended Data Fig. 2k-p). These data are consistent with the Warburg-like metabolism of PSM cells, wherein active glycolysis producing large amounts of lactate coexist with aerobic respiration [28, 29]. Importantly, stable isotope labeling patterns were almost identical between mouse and human PSM cells, suggesting that glucose and glutamine are broken down by metabolic pathways in the same way in both species and that only the rate of these processes differs between them. Together, these data confirmed that mouse PSM cells display faster mass-specific metabolic rates than human PSM cells.

To determine whether a higher mass-specific metabolic rate in mouse cells relative to human cells is observed in other cell types, we compared human and mouse neural progenitors differentiated *in vitro* from pluripotent stem cells. Simultaneous inhibition of the TGFβ and BMP pathways [30] led to the induction of PAX6+ neural progenitors after 5 days of treatment for mouse and 7 days for human PSCs (Extended Data Fig. 3a-bf). We did not detect a significant difference in volume between mouse and human PSCs (Extended Data Fig. 3c), indicating that it is not generally true that human cells are larger than their mouse equivalents. However, the mass-specific OCR and extracellular acidification rate (ECAR) were significantly higher in mouse cells than in human cells (Extended Data Fig. 3d-ei). Thus mass-specific metabolic rates are higher in different mouse cell types compared to human.

Next, we performed the mitochondrial stress test on the Seahorse instrument to measure more detailed aspects of cellular respiration beyond the basal rate. Mouse and human PSM cells differed in their maximal respiration rate and their spare respiratory capacity, both of which were significantly higher in mouse cells (Extended Data Fig. 3f-g). Respiration was less coupled to ATP production in mouse cells than human cells, thus reflecting a higher level of proton leak in mouse (Extended Data Fig. 3f,h) [31]. One potential explanation for the increased basal and maximal respiration rates of mouse cells could be a higher mitochondrial density. Indeed, comparison of the mitochondrial content between mouse and human cells using the MitoTracker Deep Red dye revealed that mouse cells contain a moderately but significantly higher number of mitochondria per unit volume (Fig. 2h). The total mitochondrial content per cell was nonetheless approximately 60% higher in human cells (Extended Data Fig. 3i), so that the oxygen consumption per mitochondrion was elevated in mouse cells relative to human cells (Extended data Fig. 3j). This was suggestive of inherent differences in mitochondrial properties between these species [32]. We thus measured the inner mitochondrial membrane potential (ΔΨm) with the dye tetramethylrhodamine methyl ester (TMRM), which displays increased fluorescence intensity at higher ΔΨm values. The TMRM signal was normalized by MitoTracker signal to account for differences in mitochondrial content between the species. This experiment revealed that human cells display a slightly higher TMRM/MitoTracker signal (Fig. 2i), suggesting that the higher OCR of mouse compared to human cells is not linked to a higher ΔΨm. The abundance and functional properties of mitochondria thus differed significantly between mouse and human PSM cells.

We next used oscillations of the segmentation clock as a proxy to investigate which aspect of metabolism controls embryonic time. In the chicken embryo, inhibition of respiration but not glycolysis alters segmentation clock oscillations [28]. We thus partially impaired the electron transport chain (ETC) with a set of small molecule inhibitors at sub-lethal concentrations in human PSM cells and measured the effect on the period of the segmentation clock. Inhibitors of the ETC complexes I (rotenone), III (antimycin a) and IV (sodium azide), which pump protons across the inner mitochondrial membrane and transfer electrons (Fig. 3a), all resulted in a significant lengthening of the segmentation clock period (Fig. 3b). Surprisingly, inhibiting the ATP synthase with oligomycin did not alter the oscillatory period (Fig. 3a-b). All these treatments resulted in reduced OCR and caused dampened oscillatory dynamics with a premature arrest of oscillations, suggesting that the segmentation clock cannot continue operating long-term when oxidative phosphorylation is impaired (Extended Data Fig. 4a-e). As ETC impairment leads to cell cycle arrest [33], we could not measure the cell cycle length under these conditions. Uncoupling the ETC from ATP production by treatment with the ionophore FCCP did not change the period either (Fig. 3b; Extended Data Fig. 4f-g), suggesting that it is the activity of the ETC rather than oxidative phosphorylation that is relevant to the segmentation clock period. These results are consistent with the phenotype of ETC mutants in *Caenorhabditis elegans*, where mutants with impaired complex I-IV function display slow larval growth, whereas ATP synthase mutants grow at normal rates [34]. It thus seems that complexes I-IV of the ETC play a conserved role in regulating developmental rate.

**Figure 3.**
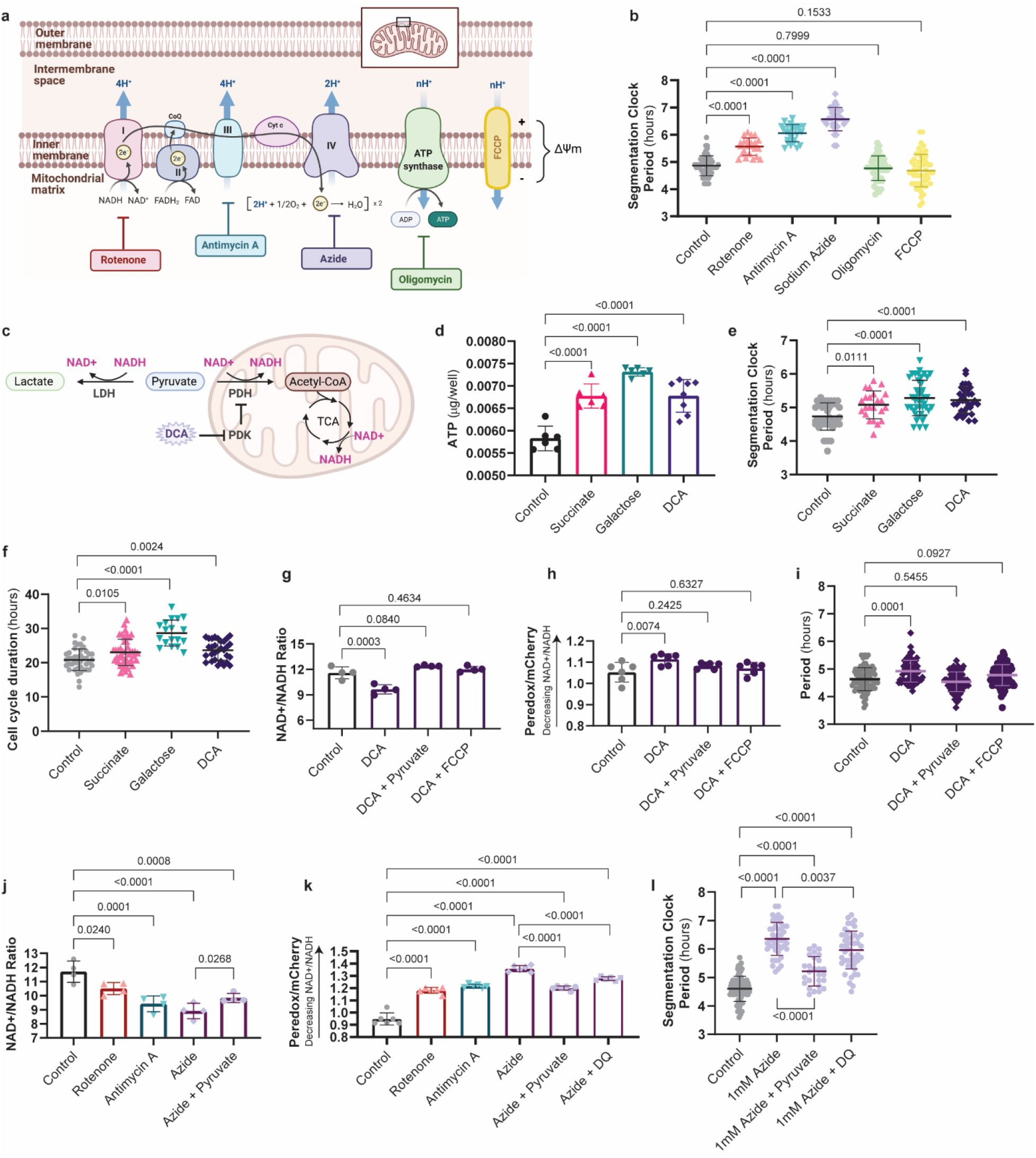
Manipulation of cellular metabolism slows down the segmentation clock by lowering the NAD^+^/NADH ratio a. Schematic drawing showing the electron transport chain and relevant small molecule inhibitors. Adapted from “Electron Transport Chain”, by BioRender.com (2021). Retrieved from https://app.biorender.com/biorender-templates b. HES7-Achilles oscillatory period in human PSM cells treated with vehicle control (DMSO; n=53), 20nM rotenone (n=23), 100nM antimycin A (n=26), 1mM sodium azide (n=30), 1μM oligomycin (n=44), and 1μM FCCP (n=55). Mean ±SD. c. Illustration depicting the alternate fates of pyruvate and their regulation by metabolic enzymes. Lactate dehydrogenase (LDH) converts pyruvate to lactate and regenerates NAD^+^ from NADH. Pyruvate dehydrogenase (PDH) oxidizes pyruvate to acetyl coenzyme-A (acetyl-CoA) in the mitochondria and consumes NAD^+^. Acetyl-CoA then enters the tricarboxylic acid (TCA) cycle, which also consumes NAD^+^. Pyruvate dehydrogenase kinase (PDK) inhibits PDH by phosphorylating it. DCA is a PDK inhibitor that promotes the conversion of pyruvate to acetyl-CoA by relieving PDH inhibition. Created with BioRender.com. d. ATP content per well for human PSM cells in control (DMSO; n=6), 25mM succinate supplementation (n=6), 10mM galactose in the absence of glucose (n=6), and 6.25mM DCA (n=8) conditions after 24 hours of culture. In each case, 30,000 cells were seeded per assay well. Each datapoint represents the average of 3 technical replicates. Mean ±SD. e. HES7-Achilles oscillatory period in human PSM cells treated with vehicle control (DMSO; n=36), 25mM succinate (n=23), 10mM galactose in the absence of glucose (n=43), and 6.25mM DCA (n=34). Mean ±SD. f. Duration of the cell cycle in hours for human PSM cells treated with vehicle control (DMSO; n=42), 25mM succinate (n=44), 10mM galactose in the absence of glucose (n=18), and 6.25mM DCA (n=30). Mean ±SD. g. Whole-cell NAD^+^/NADH ratio in vehicle-treated human PSM cells and cells treated with either 6.25mM DCA alone or DCA in combination with 1mM sodium pyruvate or 10nM FCCP for 24 hours. Each datapoint represents the average of 3 technical replicates. Mean ±SD. n=4. h. Ratiometric Peredox/mCherry signal in vehicle-treated human PSM cells and cells treated with either 6.25mM DCA alone or DCA in combination with 1mM sodium pyruvate or 10nM FCCP for 24 hours. Each datapoint represents the average of >200 individual cells analyzed within a biological replicate. Mean ±SD. n=6. i. HES7-Achilles oscillatory period in human PSM cells treated with vehicle control (DMSO; n=78), 6.25mM DCA alone (n=68), DCA with 1mM sodium pyruvate (n=73), and DCA with 10nM FCCP (n=85). Mean ±SD. j. Whole-cell NAD^+^/NADH ratio in human PSM cells treated with vehicle control (DMSO), 20nM rotenone, 100nM antimycin A, 1mM sodium azide, and 1mM sodium azide supplemented with 1mM sodium pyruvate for 24 hours. Each datapoint represents the average of 3 technical replicates. Mean ±SD. n=4. k. Ratiometric Peredox/mCherry signal in DMSO-treated human PSM cells and cells treated with 20nM rotenone, 100nM antimycin A, 1mM sodium azide alone, azide with 1mM sodium pyruvate, and azide with 5μM duroquinone for 24 hours. Each datapoint represents the average of >200 individual cells analyzed in a biological replicate. Mean ±SD. n=6. l. HES7-Achilles oscillatory period in human PSM cells treated with vehicle control (DMSO; n=67),1mM sodium azide alone (n=46), azide with 1mM sodium pyruvate (n=27), and azide with 5μM duroquinone (n=46). Mean ±SD.

The observation that ATP synthase inhibition did not significantly change the segmentation clock period suggested that cellular ATP levels are not involved in its regulation. In fact, the ADP/ATP ratio was higher in mouse PSM cells than in human cells (Fig. 2j), suggesting that an increased cellular energy charge does not mediate the accelerated developmental rate of mouse cells. To further test this hypothesis, we cultured human PSM cells under conditions that increased the cellular ATP concentration. First, we supplemented the media with succinate to directly feed the ETC at complex II. Second, we replaced glucose by galactose in the medium to increase reliance on oxidative phosphorylation for ATP production. Third, we treated the cells with the pyruvate dehydrogenase kinase (PDK) inhibitor dichloroacetate (DCA), which increases the conversion of pyruvate to acetyl-CoA for consumption by the TCA cycle (Fig. 3c). Under all three conditions, ATP levels were increased (Fig. 3d) and the OCR/ECAR ratio was increased (Extended Data Fig. 5a). We then measured the segmentation clock period and found that not only was the period not shortened, but it was moderately lengthened (Fig. 3e; Extended Data Fig. 5b-d). Under these conditions, the cell cycle length was also increased, reflecting slower proliferation (Fig. 3f). Together, these results confirmed that higher ATP concentrations do not promote faster oscillations or proliferation.

In cancer cells, PDK inhibition also diminishes the cellular proliferation rate [35]. Upon PDK inhibition, pyruvate is shunted away from lactate dehydrogenase (LDH), leading to a lower NAD^+^/NADH ratio (Fig. 3c). At the same time, complex I of the ETC cannot sufficiently regenerate cellular NAD^+^ because the inner mitochondrial membrane becomes hyperpolarized and opposes the pumping of additional protons. Supplementing cells with pyruvate, which can be rapidly reduced by LDH to generate NAD^+^, has been shown to rescue the proliferation rate of cells treated with a PDK inhibitor by restoring the NAD^+^/NADH ratio [35]. Similarly, restoring ETC activity by dissipating the elevated ΔΨm with uncoupling agents like FCCP has also been shown to rescue the NAD^+^/NADH ratio and proliferation rates [35]. We reasoned that a similar mechanism may be responsible for the increased segmentation clock period in PSM cells subjected to PDK inhibition. Indeed, we observed a significantly lower total NAD^+^/NADH ratio (Fig. 3g) and increased ΔΨm (Extended Data Fig. 5e) in PSM cells treated with DCA. The total NAD^+^/NADH ratio could be restored to control levels by pyruvate supplementation or FCCP treatment (Fig. 3g). These total NAD^+^/NADH ratio measurements included both the mitochondrial and cytoplasmic NAD(H) pools. We generated a human PSC line carrying the fluorescent sensor Peredox, which displays higher fluorescence when cytoplasmic -but not mitochondrial- levels of NADH are increased [36] (Extended Data Fig. 5f-g). Using this reporter line, we could confirm the changes in cytosolic NAD^+^/NADH ratio upon DCA treatment and its restoration by pyruvate and FCCP (Fig. 3h). Importantly, the period of the segmentation clock in cells treated with DCA was fully rescued by both pyruvate and FCCP (Fig. 3i). Together, these data suggest that the segmentation clock period depends on NAD+ availability rather than ATP supply.

Impaired ETC activity also decreases the NAD^+^/NADH ratio because the regeneration of NAD^+^ by complex I is altered. The complex I, III and IV inhibitors that cause slow segmentation clock oscillations also cause low total NAD^+^/NADH ratio and increased Peredox fluorescence (Fig. 3j-k). Pyruvate supplementation of cells treated with the complex IV inhibitor sodium azide partially restored the NAD^+^/NADH ratio [37] (Fig. 3j-k) and also partially rescued the segmentation clock period (Fig. 3l; Extended Data Fig. 5h). As an orthogonal approach, we supplemented azide-treated cells with duroquinone, which can mediate NAD^+^ regeneration from NADH via the quinone oxidase NQO1 [38]. Duroquinone treatment led to a modest recovery of the NAD^+^/NADH ratio (Fig. 3k) and slightly but significantly accelerated segmentation clock oscillations relative to cells treated with azide alone (Fig. 3l; Extended Data Fig. 5i). This suggests that ETC inhibition slowed the segmentation clock by impairing NAD redox homeostasis.

The cytoplasmic NAD^+^/NADH ratio can be manipulated by modifying the amount of pyruvate and lactate in the cell culture medium, as these substrates will shift the balance of the reaction catalyzed by LDH towards NAD^+^ or NADH production, respectively. In the presence of high glucose, pyruvate or lactate supplementation did not change the NAD^+^/NADH ratio nor did it impact the segmentation clock period (Extended Data Fig. 6a-d). Under conditions of glucose and glutamine starvation, the period of the segmentation clock was severely increased (Extended Data Fig. 6c, 6e). However, starved cells supplemented with pyruvate exhibited a higher NAD+/NADH ratio and oscillated faster than those supplemented with lactate (Extended Data Fig. 6a-c, 6e). Similarly, inhibiting LDH with sodium oxamate to prevent NAD^+^ regeneration by this enzyme led to a decreased NAD+/NADH ratio and an increased segmentation clock period (Extended Data Fig. 6f-i). Thus, the tempo of the segmentation clock could be manipulated by changing the cytosolic NAD^+^/NADH ratio.

These results suggested that differences in NAD redox balance may underlie the differences in developmental rate between mouse and human cells. NAD^+^ is used as an electron acceptor in many metabolic reactions and is required for key steps in central carbon metabolism, nucleotide synthesis, amino acid metabolism and lipid metabolism [39]. Faster developmental rates might therefore demand higher NAD^+^ availability to sustain accelerated growth and differentiation. Surprisingly, the total NAD^+^/NADH ratio was significantly lower in mouse cells compared to human cells (Fig. 2k). However, mouse PSM cells contained overall higher levels of both NAD^+^ and NADH than human cells (Extended Data Fig. 3k), suggesting that NAD^+^ and NADH concentrations may be more important than the NAD^+^/NADH ratio *per se* for the regulation of oscillatory period between species. The related dinucleotide, NADP(H), plays a central role in the defense against oxidative stress and serves as an electron donor for the synthesis of nucleotides, lipids and amino acids [40]. We therefore measured the total NADPH/NADP^+^ ratio in mouse and human PSM cells but found that it did not differ significantly between the species (Extended Data Fig. 3l).

We next asked whether these differences in respiration between mouse and human PSM cells could be responsible for the differences in protein production and degradation rates that were suggested to underlie species-specific developmental timing [1, 2]. Indeed, increased mitochondrial activity is correlated with faster transcription, faster translation, and accelerated growth in human cell lines [41–43]. We first compared translation rates by pulsing PSM cells of both species with puromycin for one hour. We subsequently detected the amount of puromycilated peptides produced during this time, a surrogate measure of the rate of total protein translation, using either antibodies or click chemistry (Fig. 4a). These experiments indicated that the mass-specific translation rate is almost twice as fast in mouse PSM cells compared to human cells and therefore scales with developmental rate (Fig. 4b; Extended Data Fig. 7a). We next showed that globally slowing down translation by treating human PSM cells with low doses of the elongation inhibitor cycloheximide results in increased clock and cell cycle periods (Fig. 4c-d; Extended Data Fig. 7b-c). This suggests that the global translation rate could potentially control developmental rate.

**Figure 4.**
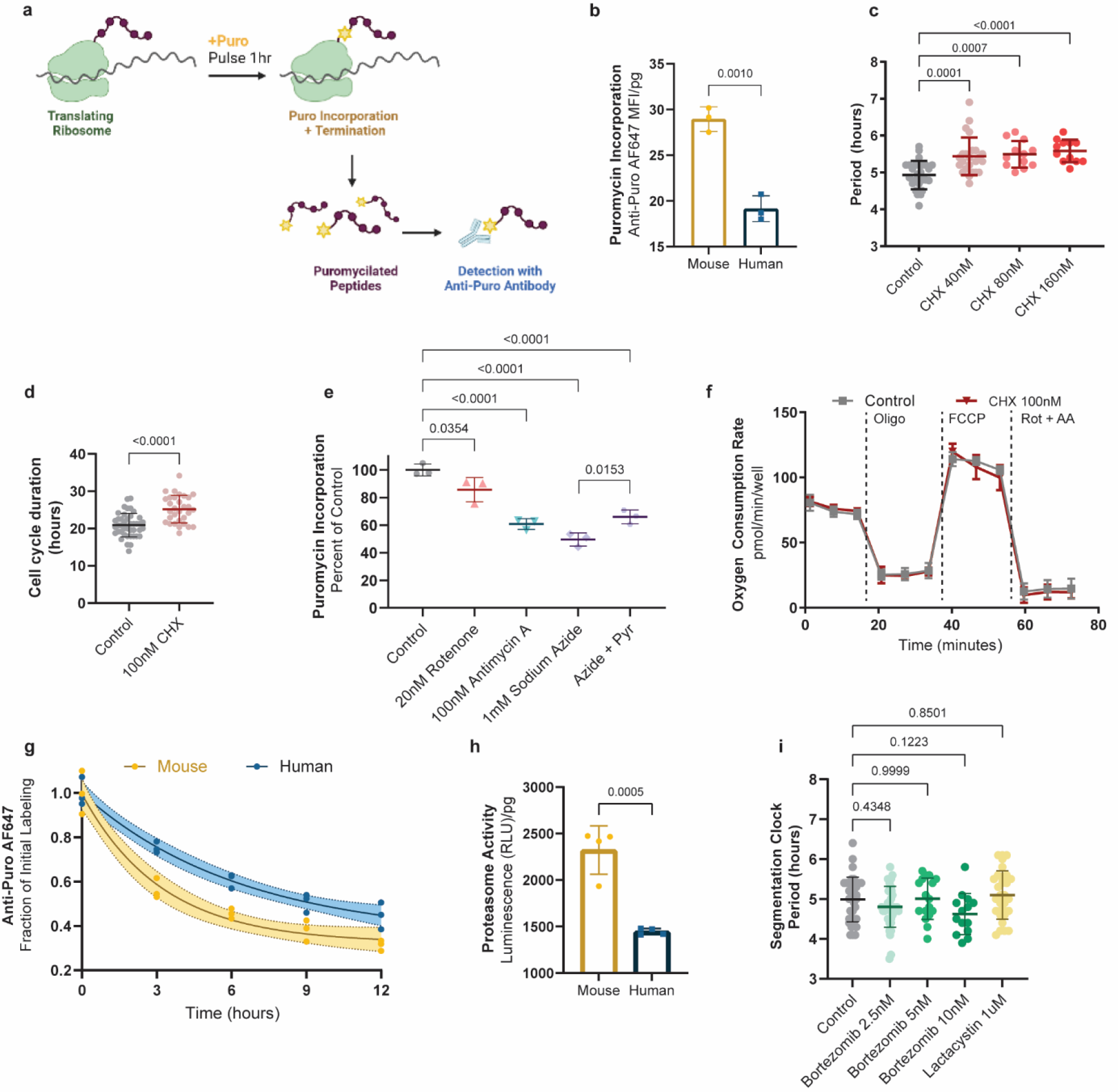
The global rate of protein synthesis acts downstream of the electron transport chain to regulate developmental speed a. Experimental approach to measure global protein synthesis by detection of puromycilated peptides following a 1-hour pulse with puromycin (puro). Created with BioRender.com. b. Global translation rate as measured by puromycin incorporation in MSGN1-Venus+ PSC-derived mouse and human PSM cells immediately after a 1-hour puromycin pulse and detection by directly conjugated AlexaFluor647 anti-puromycin antibody. Mean ±SD. n=3. c. Period of HES7-Achilles oscillations in human PSM cells treated with vehicle control (DMSO, n=27) or increasing doses of cycloheximide (40nM, n=24; 80nM, n=12; 160nM, n=12). Mean ±SD. d. Duration of the cell cycle in control (DMSO-treated; n=42) human PSM cells and cells treated with 100nM cycloheximide (CHX; n=31). Mean ±SD. e. Relative translation rate expressed as puromycin incorporation normalized to control (DMSO treatment) in human PSM cells treated with 20nM rotenone, 100nM antimycin A, 1mM sodium azide, and azide with 1mM sodium pyruvate for 24 hours. Mean ±SD. n=3. f. Oxygen consumption rate measured over the course of the mitochondrial stress test for human PSM cells treated with DMSO control or 100nM cycloheximide for 24 hours. Oligomycin, FCCP, and Rotenone + Antimycin A were added at the timepoints marked by dotted lines. Mean ±SD. n=9. g. Pulse-chase experiment tracking the degradation of puromycilated peptides over the course of 12 hours in MSGN1-Venus+ PSC-derived mouse and human PSM cells following a 1-hour pulse with puromycin. Solid line represents best one-phase decay fit with the 95% confidence intervals shown as shaded regions. n=3. h. Mass-specific proteasome activity in MSGN1-Venus+ PSC-derived mouse and human PSM cells as measured by cleavage of a luminogenic proteasome substrate. Mean ±SD. n=4. i. Period of HES7-Achilles oscillations in human PSM cells treated with DMSO control (n=35), 2.5nM (n=37), 5nM (n=17) or 10nM (n=14) bortezomib, or 1μM lactacystin (n=30). Mean ±SD.

Next, we asked if the ETC inhibitors that slow down the segmentation clock can impact the translation rate. Indeed, rotenone, antimycin A and sodium azide all decreased the translation rate as measured by puromycin incorporation (Fig. 4e). Strikingly, the magnitude of the effect on the segmentation clock period scaled with the magnitude of translation rate reduction for these inhibitors. Supplementing azide-treated cultures with pyruvate to partially restore the NAD^+^/NADH ratio also partially rescued the translation rate (Fig. 4e). Cultures treated with sodium azide for only one hour already displayed reduced translation, indicating rapid downregulation of protein synthesis upon ETC inhibition (Extended Data Fig. 7d). These results suggested that ETC inhibition and NAD^+^ depletion slow down the segmentation clock at least in part by reducing the protein production rate. To confirm this epistatic relationship, we performed Seahorse experiments in human PSM cells treated with cycloheximide. All aspects of respiration, including the basal and maximal rates, the proton leak and the non-mitochondrial respiration, were indistinguishable between cycloheximide-treated and control cells (Fig. 4f; Extended Data Fig. 7e-f). We thus concluded that mitochondrial activity acts upstream of the translation rate to control the segmentation clock period.

Differences in global protein stability also correlate with species-specific developmental rates, as the proteome half-life is 2-fold shorter in mouse neural progenitors compared to human [2]. In PSM cells, the half-life of puromycilated peptides was indeed shorter in mouse PSM cells compared to human cells (Fig. 4g). Given that the proteasome is responsible for the degradation of most cellular proteins, we compared the activity of the proteasome between the two species. As expected, the mass-specific proteasome activity was significantly elevated in mouse PSM cells (Fig. 4h), suggesting that the reduced protein stability observed in this species is caused by higher proteasome activity.

We next tested whether manipulating global protein stability could affect the segmentation clock period. High levels of proteasome inhibition led to an arrest of segmentation clock oscillations as negative feedback loops are disrupted [44]. Treating human PSM cells with low doses of the proteasome inhibitors bortezomib or lactacystin caused premature arrest of oscillations but did not change the oscillatory period despite significant inhibition of proteasome activity (Fig. 4i; Extended Data Fig. 7g-h). The cell cycle duration could not be assessed under these conditions as proteasome inhibitors induce cell cycle arrest [45]. In addition, azide treatment, which slowed down oscillations, did not reduce proteasome activity or increase the stability of puromycilated peptides (Extended Data Fig. 7i-j). To assess the degradation profile of the full-length proteome, we performed pulse-chase experiments with the methionine analog L-azidohomoalanine (AHA). Unlike puromycin, AHA incorporation into growing peptides does not induce chain termination and thus labels full-length proteins. AHA labeling resulted in indistinguishable decay profiles between control and azide-treated cells over the timeframe relevant to segmentation clock oscillations (Extended Data Figure 7k). However, protein degradation seemed to stall after 18 hours of azide treatment, suggesting that a lack of protein turnover may contribute to premature arrest of oscillations upon ETC inhibition (Extended Data Figure 7k). Together, these experiments suggest that the segmentation clock period is more sensitive to the rate of protein production than the rate of protein degradation.

In summary, we found that mass-specific respiration rates scale with and regulate the segmentation clock period by modulating NAD redox balance and, more downstream, the global translation rate. Given that the segmentation clock period can be used as a proxy for developmental rate [1], our results may explain at least in part the differences in developmental rate observed at early stages of mouse and human embryonic development. Such a mechanism may also be modulated locally in the embryo to generate heterochronic changes in segmentation, as for instance in the acceleration of segmentation clock speed relative to developmental rate in snakes [46]. However, additional studies in other embryonic cell types and other mammalian species are still required to assess the generality of our findings. In particular, it remains unclear whether cells not undergoing Warburg-like metabolism would similarly rely on NAD^+^ over ATP [47]. Moreover, distinct mechanisms may control developmental speed at fetal stages of development, when the difference in both size and developmental rate quickly increases between mouse and human [9].

Our findings also suggest that mass-specific metabolic rates for embryonic cell types may scale with adult body mass as predicted by Kleiber’s law, even under uniform culture conditions [31]. Interestingly, different cell types may achieve such scaling through opposing strategies: whereas PSM cells scaled their mass, neural progenitor cells scaled their metabolism [48]. Our studies also revealed a striking parallel between the metabolic requirements of cancer cell proliferation and those of the segmentation clock [49]. In both cases, NAD+ redox homeostasis is more important than ATP availability to maintain normal growth and oscillation rates, respectively [35]. Given that PSM cells exhibit Warburg-like metabolism with high levels of aerobic glycolysis [28], these similarities further strengthen the notion that cancer cells resemble embryonic progenitors. Moreover, this connection suggests that understanding how developmental rate is regulated will have potential applications in halting tumor growth. Similarly, the finding that the segmentation clock period is sensitive to NAD^+^ levels draws a parallel between developmental rate and aging. NAD^+^ decreases progressively with age and restoring NAD^+^ levels can ameliorate aging-associated phenotypes [50]. The aging process and developmental rate may share some regulatory mechanisms, especially given that lifespan and gestation period are positively correlated [51]. Future work should focus on the identification of factors that underlie increased mass-specific respiration rates in fast-developing species [31]. Ultimately, interspecific differences in developmental rate must be traceable to genetic causes. Lastly, the implication of translation rate downstream of mitochondrial activity strongly suggests that the integrated stress response may play a role in determining developmental rate [47, 52]. Continued research in this area will reveal how developmental time can be manipulated externally, with important applications in human stem cell therapy and *in vitro* disease modeling.

## Acknowledgements

We thank members of the Pourquié laboratory and Dr. Jason Locasale, Dr. Gary Ruvkun, Dr. Clifford Tabin, Dr. Norbert Perrimon, Dr. Vamsi Mootha, Dr. Owen Skinner and Dr. Sneha Rath for critical reading of the manuscript and discussions. The Pourquié laboratory and M.D.C. were funded by grants from the Eunice Kennedy Shriver National Institute of Child Health & Human Development (NICHD) of the National Institutes of Health under award numbers 2R01HD085121-06A and F31HD100033, respectively. This work was supported by the Cancer Systems Biology Consortium funding (U54-CA217377) from the National Cancer Institute (S.R.M.) and the Virginia and D.K. Ludwig Fund for Cancer Research (S.R.M.). The content is solely the responsibility of the authors and does not necessarily represent the official views of the National Institutes of Health. We thank the NeuroTechnology Studio at Brigham and Women’s Hospital for providing access to the Agilent Seahorse XF96 extracellular flux analyzer and Zeiss LSM880 confocal microscope, and for consultation on data acquisition and data analysis. We also thank the Center for Neurologic Diseases (ARCND) Flow Cytometry Core Facility at Brigham and Women’s Hospital for access to the BD Fortessa flow cytometer. In addition, we thank the Seahorse core facility at Brigham and Women’s hospital for training and access to the Agilent Seahorse XFe24 extracellular flux analyzer. We also express our gratitude towards Dr. Seungeun Oh and the Kirschner laboratory at Harvard Medical School for generously providing access to the Moxi Go II for cell volume measurements.

## Author Contributions

MDC and OP conceptualized the study. MDC performed the experiments and analyzed the data. TM and SM carried out suspended microchannel resonator experiments and analyzed the resulting data. DS generated the AAVS1-CAG-Peredox-mCherryNLS human PSC line. CMDG and GY contributed to the functional validation of the AAVS1-CAG-Peredox-mCherryNLS human PSC line. SG helped maintain mouse and human PSC cultures and seeded cells for differentiation. AH performed experiments involving primary mouse PSM tissue. WO helped with seahorse experiments, performed stable isotope metabolic tracing, and provided guidance on the project. MDC and OP wrote the manuscript. OP supervised the project. All authors discussed the results and commented on the manuscript.

---

**Extended Data Figure 1.**
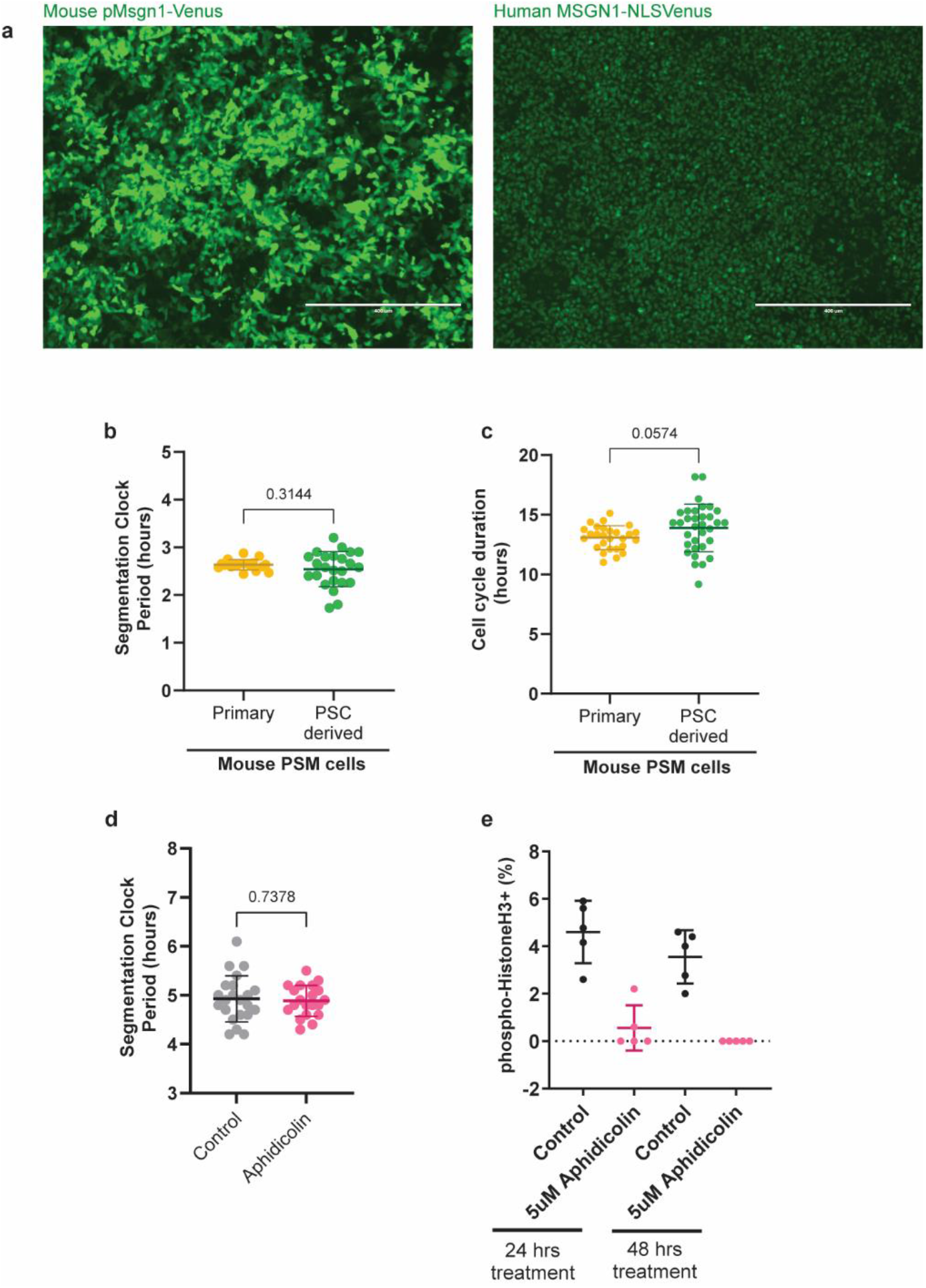
Related to Figure 1 a. MSGN1-Venus fluorescence in mouse (left) and human (right) PSC-derived PSM cells on day 2 of differentiation. Note the reporter is cytoplasmic in mouse cells but nuclear in human cells. Scale bar = 400μm. b. Period of segmentation clock oscillations in primary PSM tissue dissected from E9.5 mouse embryos carrying the *LuVeLu* reporter (n=18) and PSC-derived PSM expressing the *Hes7-Achilles* reporter (n=24). Mean ±SD. c. Cell cycle duration in primary PSM tissue dissected from E9.5 mouse embryos (n=27) and PSC-derived PSM (n=33). Mean ±SD. d. HES7-Achilles oscillatory period in human PSM cells under control (DMSO; n=22) or 5μM aphidicolin (n=20) conditions. Cultures were pre-treated with DMSO or aphidicolin for 24 hours to induce cell cycle arrest. Mean ±SD. e. Quantification of immunofluorescence staining for histone H3 phosphorylated at Ser10 in human PSM cells treated with vehicle control (DMSO) or 5 μM aphidicolin for 24 or 48 hours. Mean ±SD. n=5.

**Extended Data Figure 2.**
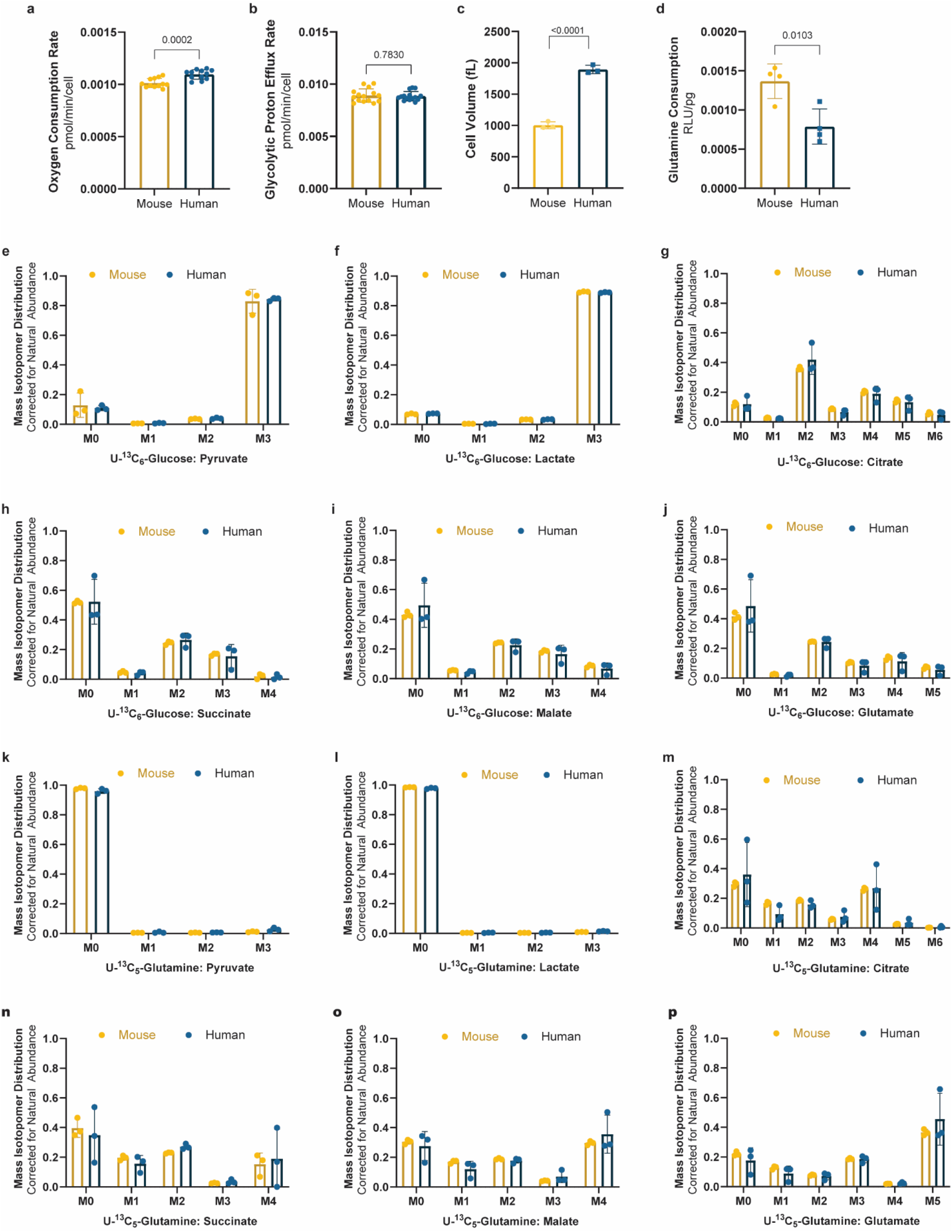
Related to Figure 2 a. Oxygen consumption rate per cell for MSGN1-Venus+ PSC-derived mouse and human PSM cells. Same data as Fig. 2d but normalized by cell. Mean ±SD. n=12. b. Glycolytic proton efflux rate per cell for MSGN1-Venus+ PSC-derived mouse and human PSM cells. Same data as Fig. 2d but normalized by cell. Mean ±SD. n=15. c. Cell volume as measured in a suspended microchannel resonator for MSGN1-Venus+ PSC-derived mouse and human PSM cells. Mean ±SD. n=3. d. Relative mass-specific glutamine consumption after 12 hours of culture for MSGN1-Venus+ PSC-derived mouse and human PSM cells. Mean ±SD. n=4. e-j. Mass isotopomer distribution, adjusted for natural abundance, for pyruvate (e), lactate (f), citrate (g), succinate (h), malate (i), and glutamate (j) after 24 hours of labeling with 25mM U^13^C_6_-Glucose in mouse and human PSM cells. Mean ±SD. n=3. k-p. Mass isotopomer distribution, adjusted for natural abundance, for pyruvate (k), lactate (l), citrate (m), succinate (n), malate (o), and glutamate (p) after 24 hours of labeling with 4mM U^13^C_5_-Glutamine in mouse and human PSM cells. Mean ±SD. n=3.

**Extended Data Figure 3.**
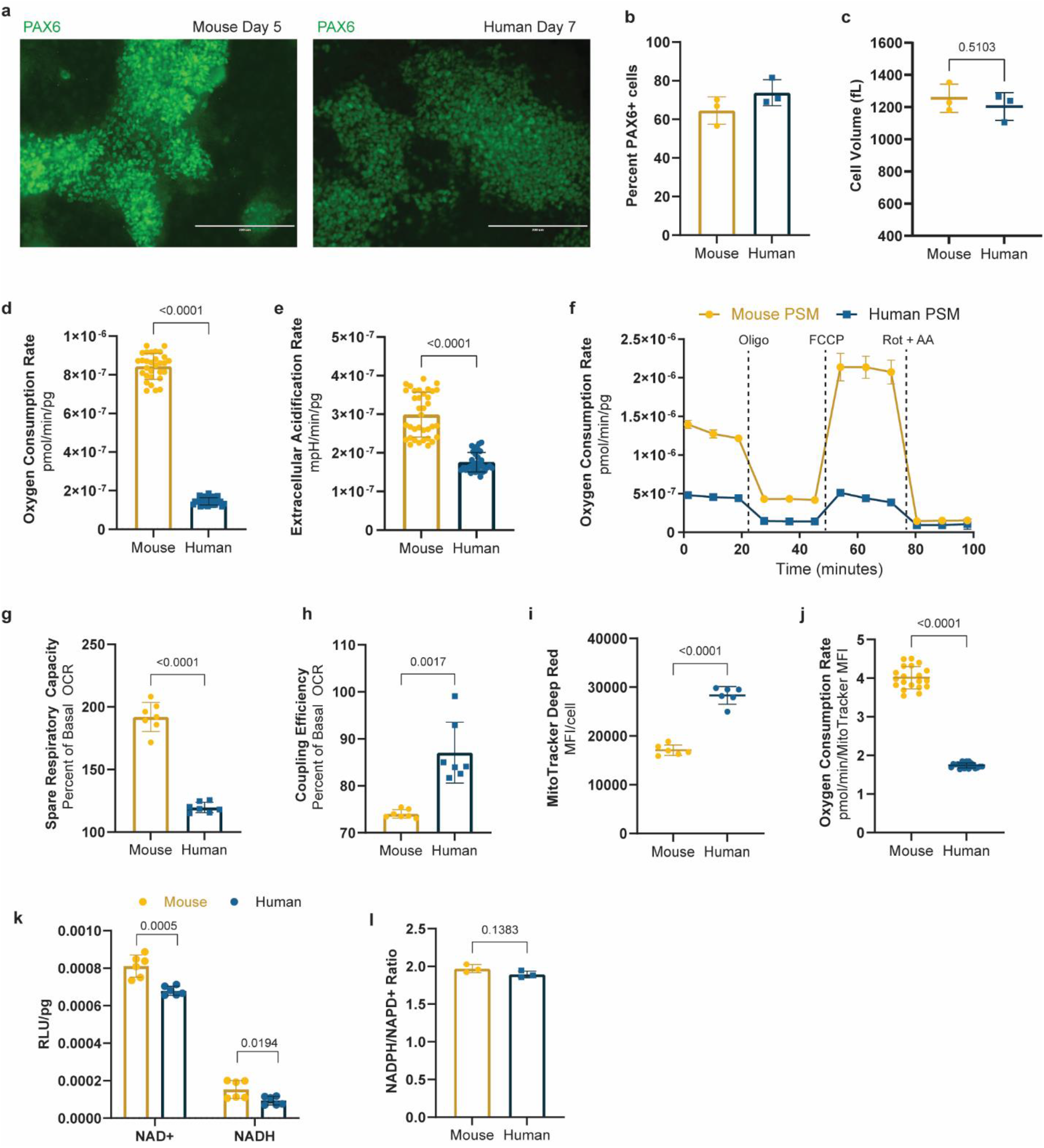
Related to Figure 2 a. Immunofluorescence staining for PAX6 in PSC-derived mouse (left) and human (right) neural progenitor cells on days 5 and 7 of differentiation, respectively. Scale bar=200μm. b. Percent PAX6+ cells in mouse (day 5) and human (day 7) neural progenitor cultures as measured by intracellular staining and flow cytometry. Mean ±SD. n=3. c. Volume of mouse (day 5) and human (day 7) neural progenitor cells as measured by a coulter counter. Mean ±SD. n=3. d. Mass-specific oxygen consumption rate for mouse (day 5) and human (day 7) neural progenitor cells. Mean ±SD. n=30. e. Mass-specific extracellular acidification rate for mouse (day 5) and human (day 7) neural progenitor cells Mean ±SD. n=36. f. Mass-specific oxygen consumption rate measured over the course of the mitochondrial stress test for MSGN1-Venus+ PSC-derived mouse (n=9) and human (n=7) PSM cells. 1μM oligomycin, 1μM FCCP, and 0.5μM Rotenone + 0.5μM Antimycin A were added at the timepoints marked by dotted lines. Mean ±SD. g. Spare respiratory capacity in MSGN1-Venus+ PSC-derived mouse and human PSM cells. Mean ±SD. n=7. h. Coupling efficiency shown as the percent of basal oxygen consumption that is linked to ATP production in MSGN1-Venus+ PSC-derived mouse and human PSM cells. Mean ±SD. n=7. i. Mitochondrial content per cell as measured by MitoTracker Deep Red staining and flow cytometry in MSGN1-Venus+ PSC-derived mouse and human PSM cells. Mean ±SD. n=6. j. Basal oxygen consumption rate normalized by mitochondrial content in MSGN1-Venus+ PSC-derived mouse and human PSM cells. Mean ±SD. n=20. k. Mass-specific NAD^+^ and NADH content in MSGN1-Venus+ PSC-derived mouse and human PSM cells. Each datapoint represents the average of 3 technical replicates. Mean ±SD. n=6. l. Whole-cell NADPH/NADP^+^ ratio in MSGN1-Venus+ PSC-derived mouse and human PSM cells. n=3.

**Extended Data Figure 4.**
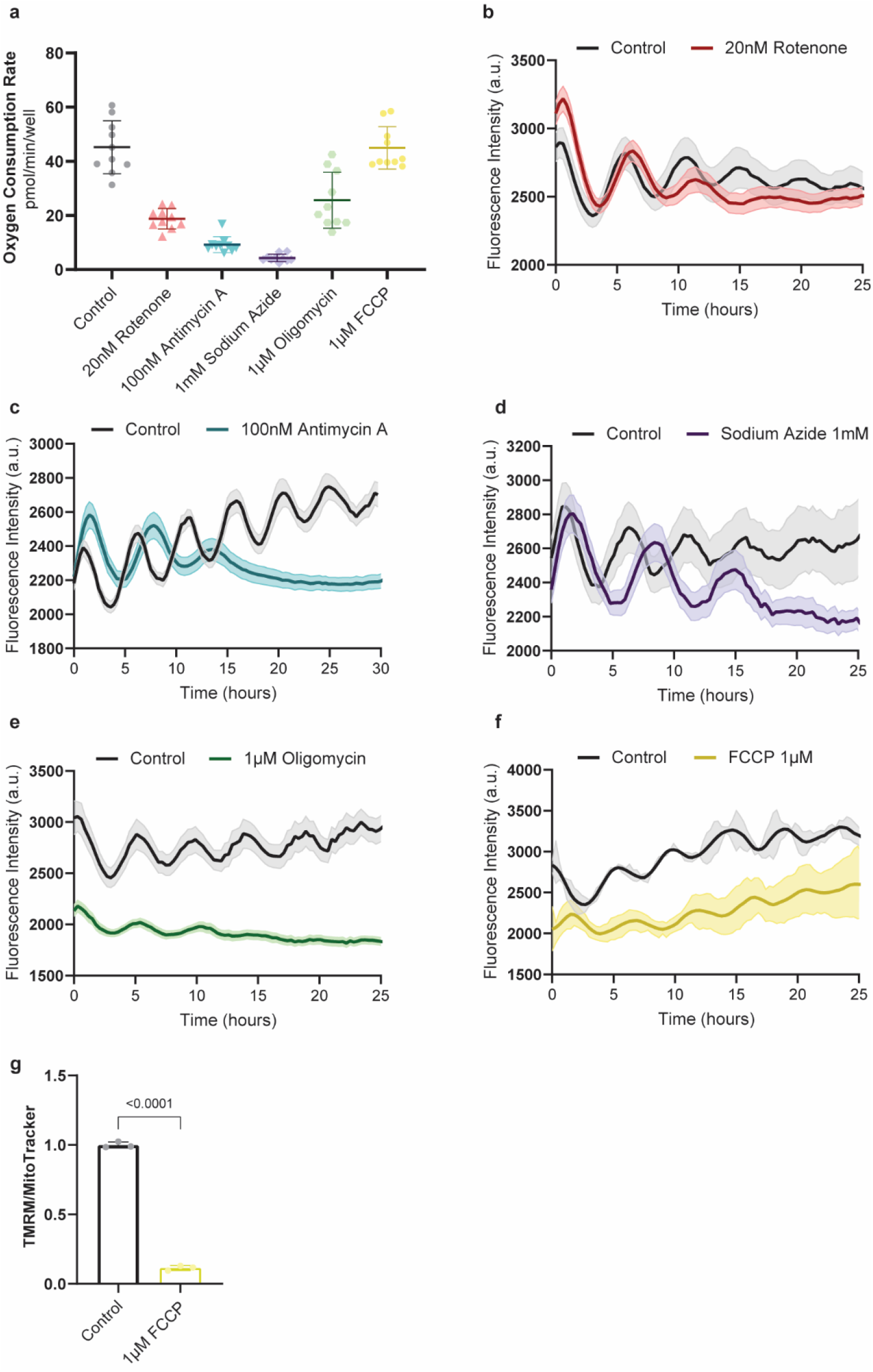
Related to Figure 3 a. Basal oxygen consumption rate in human PSM cells treated with vehicle control (DMSO), 20nM rotenone, 100nM antimycin A, 1mM sodium azide, 1μM oligomycin or 1μM FCCP. Mean ±SD. n=10. b. HES7-Achilles oscillatory profile in human PSM cultures treated with DMSO control (n=10) or 20nM rotenone (n=11). Mean ±SEM. c. HES7-Achilles oscillatory profile in human PSM cultures treated with DMSO control (n=11) or 100nM antimycin A (n=13). Mean ±SEM. d. HES7-Achilles oscillatory profile in human PSM cultures treated with DMSO control (n=9) or 1mM sodium azide (n=15). Mean ±SEM. e. HES7-Achilles oscillatory profile in human PSM cultures treated with DMSO control (n=12) or 1μM oligomycin (n=17). Mean ±SEM. f. HES7-Achilles oscillatory profile in human PSM cultures treated with DMSO control (n=3) or 1μM FCCP (n=7). Mean ±SEM. g. Inner mitochondrial membrane potential (ΔΨm) in human PSM cells under control conditions or treated acutely with 1μM FCCP. TMRM fluorescence was normalized by mitochondrial content following flow cytometry. Mean ±SD. n=3.

**Extended Data Figure 5.**
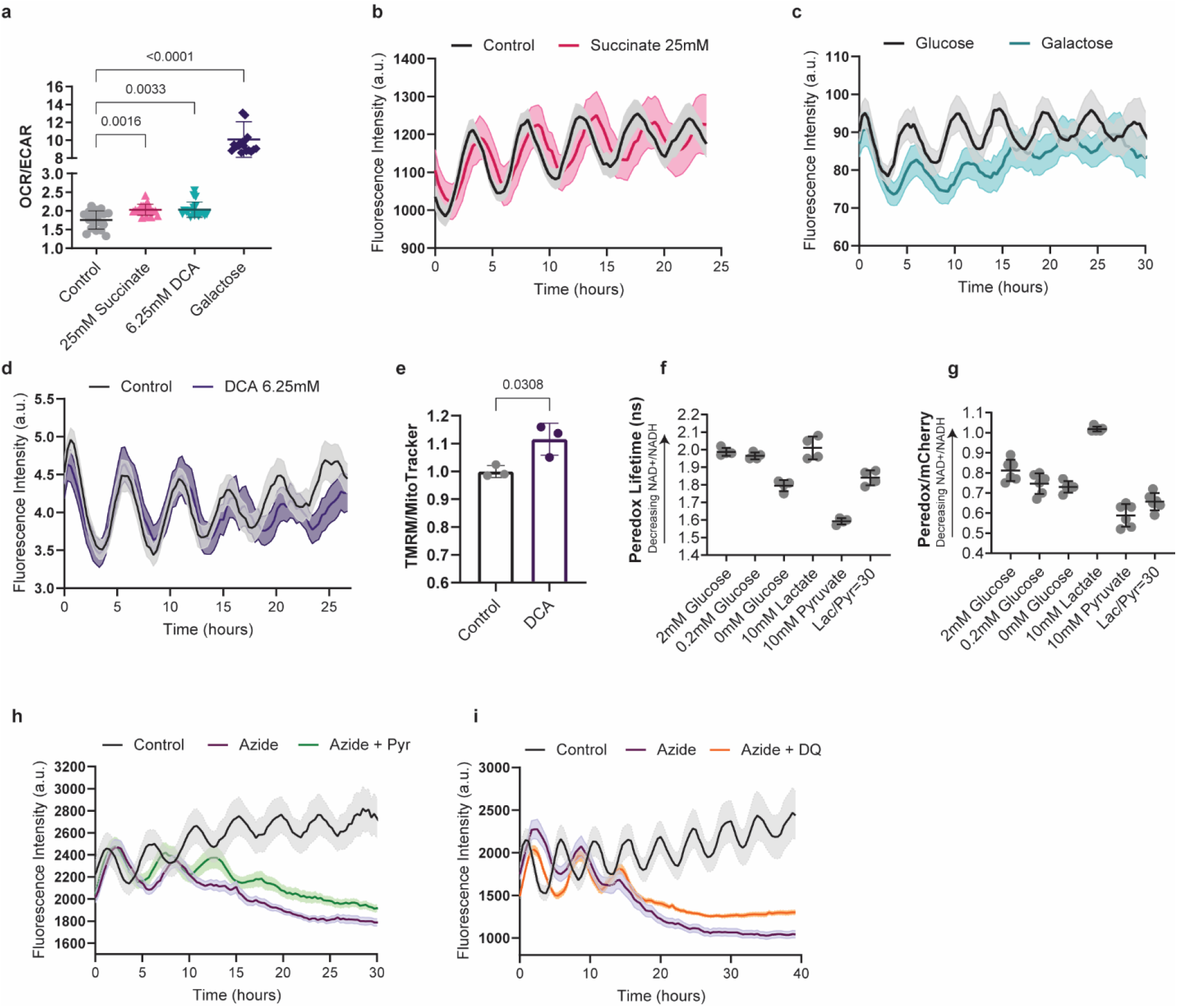
Related to Figure 3 a. Oxygen consumption rate (OCR) to extracellular acidification rate (ECAR) ratio in human PSM cells treated with DMSO control, 25mM succinate, 6.25mM DCA, or cultured with 10mM galactose instead of glucose. Mean ±SD. n=18. b. HES7-Achilles oscillatory profile in human PSM cells cultured under control conditions or supplemented with 25mM succinate. Mean ±SEM. n=8. c. HES7-Achilles oscillatory profile in human PSM cells cultured with either 10mM glucose or 10mM galactose. Mean ±SEM. n=6. d. HES7-Achilles oscillatory profile in human PSM cultures treated with vehicle control (DMSO; n=9) or 6.25mM DCA (n=8). Mean ±SEM. e. Inner mitochondrial membrane potential (ΔΨm) in human PSM cells under control conditions or treated with 6.25mM DCA for 24 hours. TMRM fluorescence was normalized by mitochondrial content following flow cytometry. Mean ±SD. n=3. f. Peredox-mCherryNLS fluorescence lifetime in human PSM cells cultured acutely in a balanced salt solution and supplemented with the indicated concentrations of glucose, lactate or pyruvate. Mean ±SD. n=4 g. Ratiometric Peredox-to-mCherry fluorescence signal in human PSM cells cultured acutely in a balanced salt solution and supplemented with the indicated concentrations of glucose, lactate or pyruvate. Mean ±SD. n=6 h. HES7-Achilles oscillatory profile in human PSM cells cultures treated with DMSO control (n=10), 1mM sodium azide alone (n=13), and azide with 1mM sodium pyruvate (n=13). Mean ±SEM. i. HES7-Achilles oscillatory profile in human PSM cells cultures treated with DMSO control (n=11), 1mM sodium azide alone (n=19), and azide with 5μM duroquinone (n=21). Mean ±SEM.

**Extended Data Figure 6.**
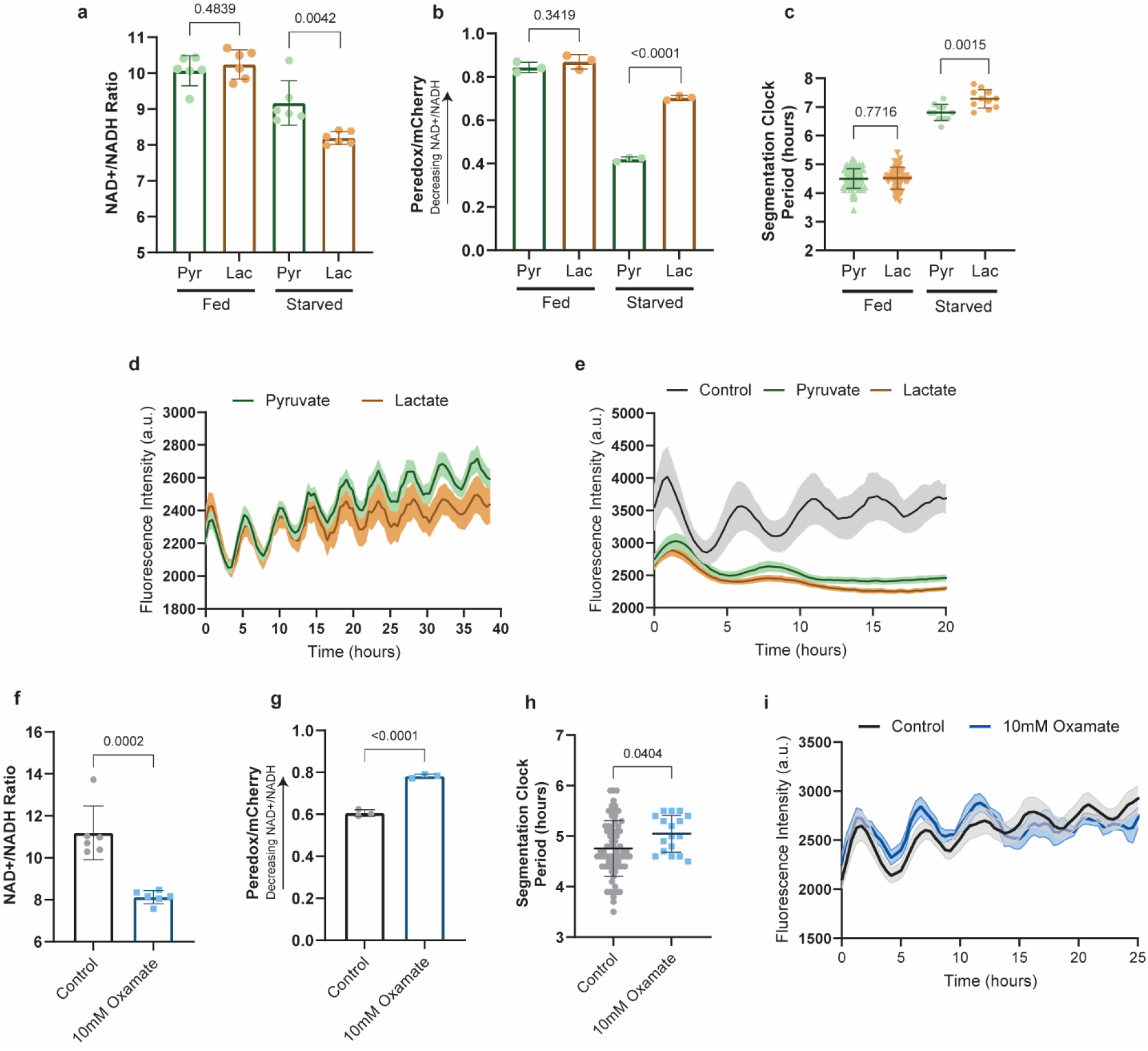
Related to Figure 3 a. Whole-cell NAD^+^/NADH ratio human PSM cells cultured under fed (17.5mM glucose, 2.5mM glutamine) or starved conditions (0mM glucose, 0mM glutamine) and supplemented with either 1mM sodium pyruvate or 1mM lactate. Each datapoint represents the average of 3 technical replicates. Mean ±SD. n=6. b. Ratiometric Peredox/mCherry signal in human PSM cells cultured under fed (17.5mM glucose, 2.5mM glutamine) or starved conditions (0mM glucose, 0mM glutamine) and supplemented with either 1mM sodium pyruvate or 1mM lactate. Each datapoint represents the average of >200 individual cells analyzed within a biological replicate. Mean ±SD. n=3. c. Period of HES7-Achilles oscillations in human PSM cells cultured under fed (17.5mM glucose, 2.5mM glutamine) or starved conditions (0mM glucose, 0mM glutamine) and supplemented with either 1mM sodium pyruvate or 1mM lactate. Mean ±SD. Fed-Pyr n=105, Fed-Lac n=64, Starved-Pyr and Starved-Lac n=11. d. HES7-Achilles oscillatory profile for human PSM cells cultured in DMEM/F12 medium containing 17.5mM glucose and 2.5mM glutamine, supplemented with either 1mM sodium pyruvate (n=14) or 1mM lactate (n=8). Mean ±SEM. e. HES7-Achilles oscillatory profile for human PSM cells cultured in the absence of glucose and glutamine, and supplemented with either 1mM sodium pyruvate or 1mM lactate. Mean ±SEM. n=11. f. Whole-cell NAD^+^/NADH ratio human PSM cells under control conditions or treated acutely with 10mM oxamate. Each datapoint represents the average of 3 technical replicates. Mean ±SD. n=6. g. Ratiometric Peredox/mCherry signal in human PSM cells cultured control conditions or treated acutely with 10mM oxamate. Each datapoint represents the average of >200 individual cells analyzed within a biological replicate. Mean ±SD. n=3. h. Period of HES7-Achilles oscillations in human PSM cells cultures treated with DMSO control (n=73) or 10mM sodium oxamate (n=17). Mean ±SEM. i. HES7-Achilles oscillatory profile in human PSM cells cultures treated with DMSO control (n=10) or 10mM sodium oxamate (n=3). Mean ±SEM.

**Extended Data Figure 7.**
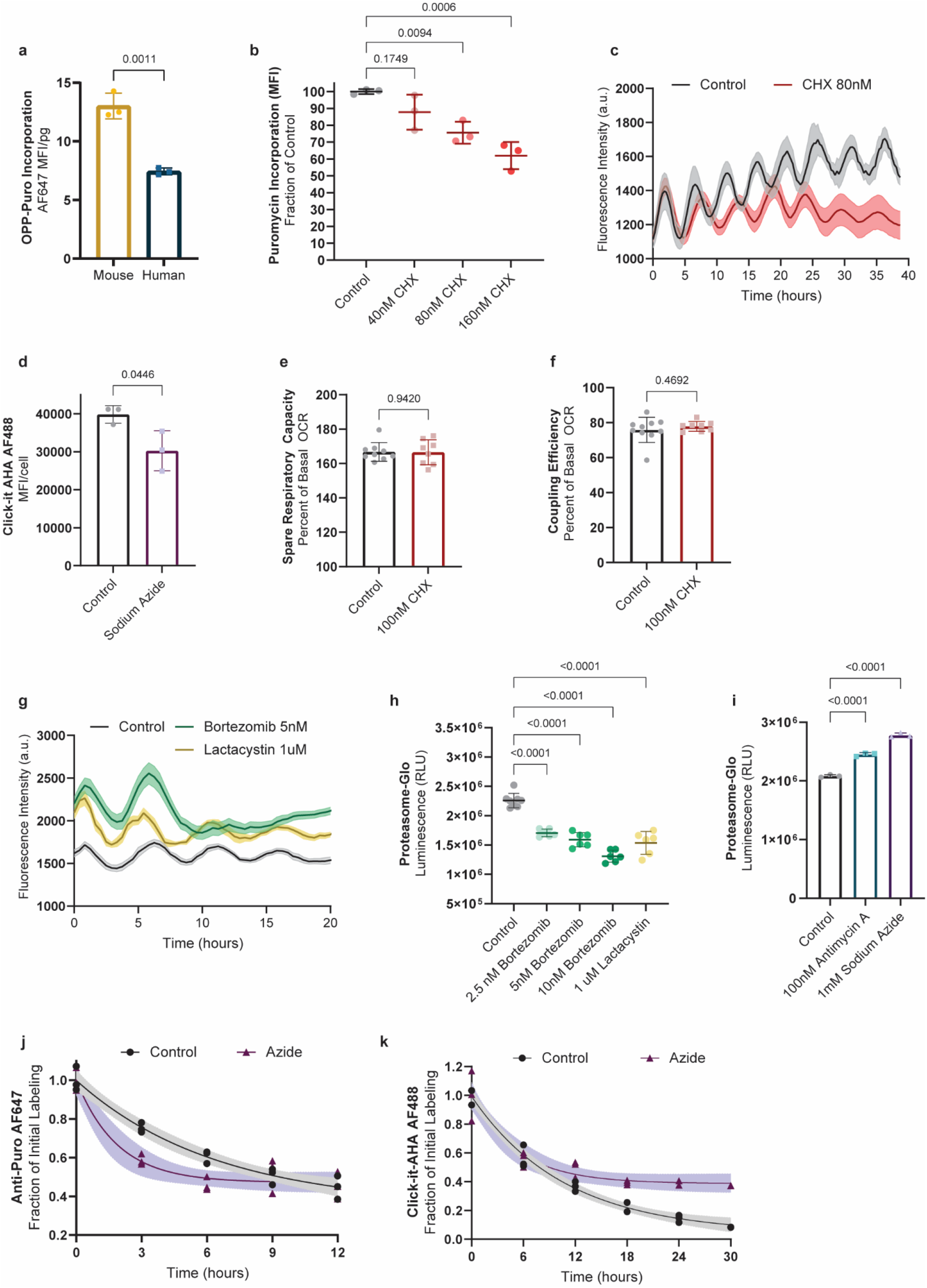
Related to Figure 4 a. OPP-Puromycin incorporation as a measure of global translation rate in MSGN1-Venus+ PSC-derived mouse and human PSM cells. OPP-Puromycilated peptides were detected by click chemistry with AlexaFluor647-Picoyl Azide. Mean ±SD. n=3. b. Translation rate as measured by puromycin incorporation expressed as percent of control for human PSM cells treated with 40nM, 80nM or 160nM cycloheximide (CHX) for 24 hours. Mean ±SD. n=3. c. HES7-Achilles oscillatory profile for human PSM cells treated with DMSO-control (n=3) or 80nM cycloheximide (CHX; n=5). Mean ±SEM. d. Translation rate as measured by incorporation of the methionine analog AHA in human PSM cells treated with either DMSO control or 1mM sodium azide for one hour. Mean ±SD. n=3. e. Spare respiratory capacity in human PSM cells treated with vehicle control (DMSO) or 100nM cycloheximide (CHX). Mean ±SD. n=8. f. Coupling efficiency shown as the percent of basal oxygen consumption that is linked to ATP production in human PSM cells treated with vehicle control (DMSO) or 100nM cycloheximide (CHX). Mean ±SD. n=8. g. HES7-Achilles oscillatory profile for human PSM cells treated with DMSO-control (n=9), 5nM bortezomib (n=13), or 5μM lactacystin (n=8). Mean ±SEM. h. Proteasome activity cells as measured by cleavage of a luminogenic proteasome substrate in human PSM treated with DMSO control, 2.5nM, 5nM or 10nM bortezomib, or 1μM lactacystin. Mean ±SD. n=6. i. Proteasome activity cells as measured by cleavage of a luminogenic proteasome substrate in human PSM treated with DMSO control, 100nM antimycin A, or 1mM sodium azide. Mean ±SD. n=3. j. Pulse-chase experiment tracking the degradation of puromycilated peptides over the course of 12 hours in human PSM cells treated with DMSO control or 1mM sodium azide following a 1-hour pulse with puromycin. Solid line represents best one-phase decay fit with the 95% confidence intervals shown as shaded regions. n=3. k. Pulse-chase experiment tracking the degradation of AHA-labeled proteins over the course of 30 hours in human PSM cells treated with DMSO control of 1mM sodium azide following a 1-hour pulse with AHA. Solid line represents best one-phase decay fit with the 95% confidence intervals shown as shaded regions. n=3

**Extended Data Table 1.**
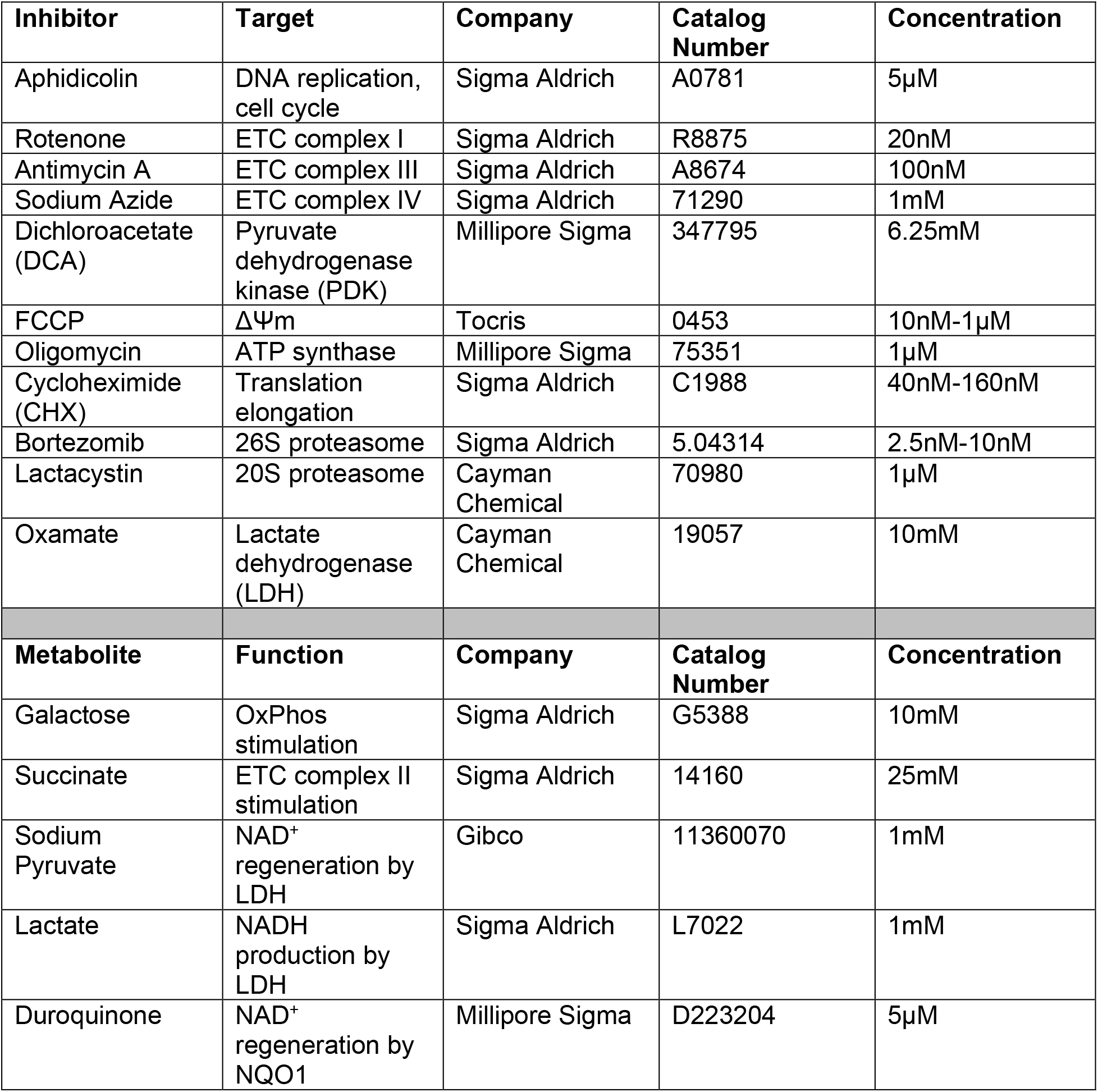
List of small molecule inhibitors and metabolite supplements

| Inhibitor | Target | Company | Catalog Number | Concentration |
| --- | --- | --- | --- | --- |
| Aphidicolin | DNA replication, cell cycle | Sigma Aldrich | A0781 | 5μM |
| Rotenone | ETC complex I | Sigma Aldrich | R8875 | 20nM |
| Antimycin A | ETC complex III | Sigma Aldrich | A8674 | 100nM |
| Sodium Azide | ETC complex IV | Sigma Aldrich | 71290 | 1mM |
| Dichloroacetate (DCA) | Pyruvate dehydrogenase kinase (PDK) | Millipore Sigma | 347795 | 6.25mM |
| FCCP | $\Delta\Psi_m$ | Tocris | 0453 | 10nM-1μM |
| Oligomycin | ATP synthase | Millipore Sigma | 75351 | 1μM |
| Cycloheximide (CHX) | Translation elongation | Sigma Aldrich | C1988 | 40nM-160nM |
| Bortezomib | 26S proteasome | Sigma Aldrich | 5.04314 | 2.5nM-10nM |
| Lactacystin | 20S proteasome | Cayman Chemical | 70980 | 1μM |
| Oxamate | Lactate dehydrogenase (LDH) | Cayman Chemical | 19057 | 10mM |
| Metabolite | Function | Company | Catalog Number | Concentration |
| Galactose | OxPhos stimulation | Sigma Aldrich | G5388 | 10mM |
| Succinate | ETC complex II stimulation | Sigma Aldrich | 14160 | 25mM |
| Sodium Pyruvate | NAD<sup>+</sup> regeneration by LDH | Gibco | 11360070 | 1mM |
| Lactate | NADH production by LDH | Sigma Aldrich | L7022 | 1mM |
| Duroquinone | NAD<sup>+</sup> regeneration by NQO1 | Millipore Sigma | D223204 | 5μM |

## Materials and Methods

### Pluripotent Stem Cell Culture

*E14* and its derivative reporter lines *pMsgn1-Venus [1]* and *Hes7-Achilles [2]* mouse ESCs were maintained under feeder-free conditions in gelatin-coated dishes (StemCell Technologies cat. no. 07903) with 2i medium composed of high glucose DMEM (Gibco cat. no. 11965-118) supplemented with 1% GlutaMAX (Gibco cat. no. 35050061), 1% Non-Essential Amino Acids (Gibco cat. no. 11140-050), 1% Sodium Pyruvate (Gibco cat. no. 11360-070), 0.01% Bovine Serum Albumin (Gibco cat. no. 15260-037), 0.1% β-mercaptoethanol (Gibco cat. no. 21985-023), 15% Fetal Bovine Serum (EMD Millipore cat. no. ES009B), 1000 U/mL LIF (EMD Millipore cat. no. ESG1106), 3μM CHIR99021 (Tocris cat. no. 4423) and 1μM PD0325901 (Stemgent cat. no. 04-006). mESCs were passaged by TrypLE Express (Gibco cat. no. 12605010) dissociation every two days at a density of 1× 10^4^ cells per cm^2^.

Human stem cell work was approved by Partners Human Research Committee (Protocol Number 2017P000438/PHS). We complied with all relevant ethical regulations. Written informed consent from the donor of the NCRM1 iPS cells was obtained by Rutgers University at the time of sample collection. *NCRM1* iPS cells (RUCDR, Rutgers University) and lines carrying the *MSGN1-Venus [3]*, *HES7-Achilles [2], HES7-Achilles; pCAG-H2B-mCherry [2]* reporters and the *AAVS1-CAG-Peredox-mCherry-NLS* sensor were maintained in Matrigel-coated plates (Corning, cat. no. 35277) in mTeSR1 medium (StemCell Technologies cat. no. 05851) as previously described [2, 4]. hiPSCs were passaged every 4-5 days by Accutase (Corning cat. no. 25058CI) dissociation and seeded at a density of 5× 10^4^ cells per cm^2^ in mTeSR1 supplemented with 10μM Y-27362 dihydrochloride (Rocki; Tocris Bioscience, cat. no. 1254).

### Presomitic Mesoderm Differentiation

Mouse ESCs were pre-differentiated to an epiblast-like state as previously described [2] by seeding fibronectin-coated dishes (BD Biosciences cat. no. 356008) at a density of 0.8× 10^4^ cells per cm^2^ in NDiff227 media (Takara cat. no. Y40002) supplemented with 1% KSR (Gibco cat. no. 10828028), 25 ng/ml Activin A (R&D systems cat. no. 338-AC-050) and 12 ng/ml bFGF (PeproTech cat. no. 450-33). The medium was refreshed after 24 hours.

Human iPSCs were seeded on Matrigel-coated plates at a density of 3× 10^4^ cells per cm^2^ in mTeSR1 with 10μM Rocki. At 24 hours, the medium was replaced by mTeSR1 without Rocki.

Presomitic mesoderm differentiation was initiated 48 hours after initial seeding for both mouse and human PSCs. Cultures were switched to DMEM/F12 GlutaMAX (Gibco cat. no. 10565042) supplemented with 1% Insulin-Transferrin-Selenium (ITS; Gibco cat. no. 41400045), 5% Fetal Bovine Serum (EMD Millipore cat. no. ES009B), 6 μM Chir 99021 (Tocris cat. no. 4423), 20ng/ml murine bFGF (PeproTech cat. no. 450-33) and 30 ng/ml Activin A (R&D systems cat. no. 338-AC-050). After 24 hours, the medium was replaced by DMEM/F12 GlutaMAX with 1% ITS, 5% FBS, 6 μM Chir 99021, 20ng/ml bFGF and 0.5 μM LDN193189 (Stemgent cat. no. 04-0074). If PSM cultures needed to be maintained for longer than 2 days, the medium was refreshed at 48 hours with DMEM/F12 GlutaMAX, 1% ITS, 5% FBS, 6 μM Chir 99021, 0.5 μM LDN193189, 50 ng/ml mFgf4 (R&D Systems cat. no. 5846-F4-025), 1 μg/ml Heparin (Sigma Aldrich cat. no. H3393-100KU), 2.5 μM BMS493 (Sigma Aldrich cat. no. B6688-5MG) and 10 μM Rocki to maintain the posterior PSM fate [2, 5].

For live imaging experiments, cells were seeded on 24 well glass-bottom plates (In Vitro Scientific cat. no. P24-1.5H-N) on day 0 and cultured in DMEM/F12 without phenol red (Gibco cat. no. 31053028). Whenever the effects of chemical inhibitors or culture conditions on the segmentation clock were tested, human *HES7-Achilles* or *HES7-Achilles; AAVS1-CAG-H2B-mCherry* cells were differentiated under serum-free conditions as previously described [2] to avoid confounding factors from FBS composition.

### Neural Progenitor Differentiation

Neural progenitor induction relied on dual Smad inhibition and was adapted from previously described protocols [6]. Mouse ESCs were seeded on fibronectin-coated dishes at a density of 1× 10^4^ cells/cm^2^ and pre-differentiated to epiblast state as described above. Human iPSCs were seeded on matrigel-coated plates at a density of 3.5× 10^4^ cells per cm^2^ in mTeSR1 with 10μM Rocki. At 24 hours, the medium was replaced by mTeSR1 without Rocki. Two days after initial seeding, both mouse and human cells were switched to NDiff227 supplemented with 1% FBS, 0.1 μM LDN193189, and 10 μM SB431542 (Selleck Chemicals cat. no. S1067). The media was refreshed daily. Mouse cells were cultured for 5 days and human cells for 7 days. Neural progenitor fate was confirmed by PAX6 immunofluorescence as described below.

### Immunofluorescence

For immunostaining of 2D cultures, cells were grown on Matrigel-coated 24 well glass-bottom plates (In Vitro Scientific cat. no. P24-1.5H-N). Cells were rinsed in PBS and fixed in a 4% paraformaldehyde solution (Electron Microscopy Sciences cat. no. 15710) for 20 minutes at room temperature, then washed 3 times with phosphate buffered saline (PBS). Samples were permeabilized by washing three times for three minutes each in Tris buffered saline (TBS) with 0.1% Tween (TBST) and blocked for one hour at room temperature in TBS-0.1% Triton-3% FBS. The primary antibody (Rabbit α PAX6 Biolegend cat. no. 901301, lotB277104 or Rabbit α pHistone H3 (Ser10) Santa Cruz cat. no. sc-8656, lot D1615) was diluted in blocking solution at 1:350 and incubated overnight at 4°C with gentle rocking. Following three TBST washes and a short 10-minute block, cells were incubated with an Alexa-Fluor conjugated secondary antibody (1:500) and Hoechst33342 (1:1000) overnight at 4°C with gentle rocking. Three final TBST washes and a PBS rinse were performed, and cells were mounted in Fluoromount G (Southern Biotech cat. no. 0100-01). Images were acquired using a Zeiss LSM780 point scanning confocal microscope with a 20X objective.

### PAX6 Intracellular staining

Samples were washed in PBS and dissociated with TrypLE. One million cells per sample were fixed with 4% formaldehyde and then permeabilized with 0.3% Triton, 0.5% BSA in PBS. Cells were washed once in 0.5% BSA. PAX6 primary antibody (Rabbit α PAX6 Biolegend cat. no. 901301, lotB277104) was diluted 1:100 in 0.5% BSA and samples were incubated for 1 hour. Following a wash in PBS, Alexa-Fluor 488 conjugated secondary antibody (1:100) was applied for 30 minutes. Samples were then washed and analyzed by flow cytometry.

### Generation, Validation and Imaging of *AAVS1-CAG-Peredox-mCherry-NLS*

*CAG-Peredox-mCherry-NLS* was inserted into the *AAVS1* safe harbor locus using the approach previously described by Oceguera-Yanez and colleagues [7]. Briefly, we cloned the *CAG-Peredox-mCherry-NLS* sequence from pcDNA3.1-Peredox-mCherry-NLS (Addgene cat. no. 32384) into the pAAVS1-P-CAG-DEST vector (Addgene cat. no. 80490) by Gibson assembly and co-transfected it along with the pXAT2 vector (Addgene cat. no. 80494) into *NCRM1* cells. Two days after transfection, we selected positive clones by supplementing mTeSR1 with puromycin (0.5 μg/mL, Sigma–Aldrich cat. no. P7255) for a total of 10 days. To enhance single cell survival, we added CloneR (StemCell Technologies cat. no. 05888) to the media during the first 2 days of selection. We obtained several positive clones and confirmed the homozygous insertion of the sensor by PCR as previously described [7].

To validate that the Peredox NADH/NAD+ sensor worked as expected in the newly generated *AAVS1-CAG-Peredox-mCherry-NLS* line, we first performed fluorescence lifetime imaging (FLIM) under different conditions (Extended Data Fig. 4f). We differentiated the sensor line to PSM fate under serum-free conditions on glass coverslips. Cells on coverslips were submerged in a recording chamber filled with a balanced salt solution (140mM NaCl, 2.5mM KCl, 10mM HEPES, 1mM MgCl2, 2mM CaCl2, and pH 7.4) at ~34°C, using a perfusion rate of 5 ml/min. Cells were sequentially perfused with the following solutions: 2mM glucose, 0.2mM, 0mM glucose, 0mM glucose with 10mM lactate, 0mM glucose with 10mM pyruvate, and finally 0mM glucose with a mixture of lactate to pyruvate at a ratio of 30:1. Cells were visualized with a Thorlabs Bergamo II microscope (Thorlabs Imaging Systems, Sterling, VA), with hybrid photodetectors R11322U-40 (Hamamatsu Photonics, Shizuoka, Japan). The objective lens used for cell visualization was an Olympus LUMPLFLN 60x/W (NA 1.0). Biosensor fluorescence was excited using light with a wavelength of 790 nm, delivered by a Chameleon Vision-S tunable Ti-Sapphire mode-locked laser (80 MHz,~75 fs; Coherent, Santa Clara, CA). Fluorescence emission light was split with an FF562-Di03 dichroic mirror and bandpass filtered for green (FF01-525/50) and red (FF01-641/75) channels (all filter optics from Semrock, Rochester, NY). Peredox emission was recorded in the green channel. The photodetector signals and laser sync signals were preamplified and then digitized at 1.25 gigasamples per second using a field programmable gate array board (PC720 with FMC125 and FMC122 modules, 4DSP, Austin, TX). Microscope control and image acquisition, as well as laboratory-built firmware and software for fluorescence lifetime determination, can be found elsewhere [8].

To further confirm that similar results could be obtained by ratiometric measurements, we imaged *AAVS1-CAG-Peredox-mCherry-NLS* cells with an LSM880 confocal microscope using a 20X/0.8 objective (Extended Data Fig. 4g) PSM cells were cultured in a balanced salt solution supplemented with 2mM, 0.2mM or 0mM glucose, 10mM lactate, 10mM pyruvate or a mixture of lactate to pyruvate at a ratio of 30:1 as described above. Images were thresholded based on mCherry-NLS fluorescence and nuclei were automatically segmented in Fiji [9]. A ratio of the mean Peredox: mCherry fluorescence intensity was calculated for each cell. All cells within an image were averaged to obtain the mean ratio for each sample. This ratiometric imaging approach was used in all experiments.

### Mouse and Human PSM Co-Culture

Mouse and human PSCs were differentiated to PSM fate separately as described above. On day 2 of differentiation, PSM cells were dissociated with TrypLE. Mixed cell suspensions consisting of *HES7-Achilles; AAVS1-CAG-H2B-mCherry* human cells combined with either human NCRM1 or mouse E14 cells at a ratio of 1:100 were prepared and reseeded on fibronectin-coated 24 well glass-bottom plates at a density of 2.5 x 10^6^ cells per cm^2^. Cells were then allowed to attach for one hour and subjected to timelapse imaging. AAVS1-CAG-H2B-mCherry+ cells were automatically segmented and tracked as previously described [2] to obtain single-cell HES7-Achilles fluorescence intensity profiles.

### Primary Mouse PSM Explant culture

Explant culture was performed as described by Hubaud and colleagues [5]. *LuVeLu* [10] CD1 E9.5 mice (both male and female) were sacrificed according to local regulations in agreement with national and international guidelines. We complied with all relevant ethical regulations. Study protocol was approved by Brigham and Women’s Hospital IACUC/CCM (Protocol number N000478). Sample size was not estimated, nor were randomization or blinding performed. Tailbud was dissected with a tungsten needle and ectoderm was removed using Accutase. Explants were then cultured on fibronectin-coated plate (LabTek chamber). The medium consisted of DMEM, 4.5g/L Glucose, 2mM L-Glutamine, non-essential amino acids 1x, Penicillin 100U/mL, Streptomycin 100μg/mL, 15% fetal bovine serum (FBS), Chir-99021 3μM, LDN193189 200nM, BMS-493 2.5 μM, mFgf4 50ng/mL, heparin 1μg/mL, HEPES 10mM and Y-27632 10μM. Explants were incubated at 37°C, 7.5% CO_2_. Live imaging was performed on a confocal microscope Zeiss LSM 780, using a 20X objective (note that the tiling could create lines between the different images).

### Timelapse Microscopy

Time lapse-imaging of mouse and human PSM cells was performed on a Zeiss LSM 780 point-scanning confocal inverted microscope fitted with a large temperature incubation chamber and a CO_2_ module. An Argon laser at 514 nm and 7.5% or 2% power was used to excite Achilles or Venus fluorescent proteins, respectively. A DPSS 561 laser at 561nm and 2% laser power was used to excite mCherry. In all cases, a 20X Plan Apo (N.A. 0.8) objective was used to acquire images with an interval of 18 or 25 minutes in the case of human samples and 12 minutes for mouse samples, for a total of 24-48 hours. A 3×3 tile of 800×800 pixels per tile with a single z-slice of 18 μm thickness and 16-bit resolution was acquired per position. Multiple positions, with at least two positions per sample, were imaged simultaneously using a motorized stage. Mouse explant imaging was performed on a Zeiss LSM780 microscope using a 20X/0.8 objective. Single section (~19.6μm thick) with tiling (3×3) of a 512×512 pixels field was acquired every 7.5 minutes at 8-bit resolution.

### Oscillation analysis

Time lapse movies of HES7-Achilles were first stitched and separated into subsets by position in the Zen program (Zeiss). Then, background subtraction and Gaussian blur filtering were performed in Fiji [9] to enhance image quality. A small region of interest (ROI) was drawn and the mean fluorescence intensity over time was calculated. Intensity is presented in arbitrary units. For figure 1e, intensity profiles were smoothened in GraphPad Prism using 6 neighboring data points and a 2^nd^ order polynomial. Profiles were then normalized between zero and one. The oscillatory period was defined as the average time between two peaks in *HES7-Achilles* profiles.

### Cell Cycle Length Measurements

To generate cultures with sparsely labeled cells, we mixed *HES7-Achilles; AAVS1-CAG-H2B-mCherry* human iPSCs or *pMsgn1-Venus* mouse ESCs with their parental line (*NCRM1* or *E14*, respectively) at a ratio of 1:100 during the initial seeding. Cultures were then differentiated normally and subjected to timelapse imaging. Individual reporter cells were tracked manually on Fiji [9] and the time of cell division was recorded. The cell cycle length was defined as the time elapsed between the time a cell first divides and the time one of its daughter cells divides again.

To measure cell cycle length in primary mouse PSM explants, lentiviral infection was used to sparsely label cells with an *SV40-mCherry* reporter [5]. The plasmid E[beta]C (Addgene cat. no. 24312) was used. Lentivirus was produced in 293T cells, which were transfected using the CaCl2 method with the packaging plasmids psPAX2 (Addgene cat. no. 12260) and pVSVG (gift from M. Wernig lab). Supernatant was collected, filtered using a 0.45μM filter and concentrated by centrifugating 4 volumes of supernatant on 1 volume of TNE buffer (50mM Tris pH7.2, 100mM NaCl, 0.5mM EDTA, 15% sucrose) at 7197 relative centrifugal field (rcf) for 4 hours at 4°C. Explants were infected for ~4 hours and further incubated overnight before imaging. *SV40-mCherry+* cells were manually tracked, and the cell cycle length was calculated as described above.

### Flow Cytometry and Cell Sorting

To determine the fraction of PSM cells *expressing MSGN1-Venus,* cultures were dissociated with Accutase and analyzed by flow cytometry using an S3 cell sorter (Biorad). Undifferentiated ESCs or iPS cells, which do not express the fluorescent protein, were used as a negative controls for gating purposes. Samples were analyzed in biological triplicates. Results are presented as the percentage of Venus-positive cells among singlets. The same gating strategy was used to sort MSGN1-Venus+ for subsequent experiments. All other flow cytometry analyses were performed on a 5-laser Fortessa analyzer (BD). Automatic compensation was set up whenever more than one stain or fluorescent protein was used at a time. Flow cytometry data was analyzed in FlowJo. The mean fluorescence intensity (MFI) for 10,000 cells is presented. In the case of mouse vs. human comparisons, only MSGN1-Venus+ cells were considered in the analysis and MFI was normalized to cell mass.

### Mitochondrial content and ΔΨm measurement

PSM cells were washed in PBS and incubated in DMEM supplemented with 25nM Mitotracker Deep Red FM (Invitrogen cat. no. M22426), 25nM Mitotracker Green (Invitrogen cat. no. M7514), and/or 20nM TMRM (Invitrogen, T668) for 30 minutes as previously described [11]. The cells were then dissociated with TrypLE, resuspended in PBS-1%FBS and analyzed by flow cytometry. As a control, 1uM FCCP was used to depolarize the inner mitochondrial membrane.

### Seahorse Assays

PSM cells were dissociated on day 2 of differentiation and reseeded onto fibronectin-coated Seahorse plates (Agilent cat. no. 101085-004) at a density of 7 x 10^5^ cells per cm^2^ in Seahorse XF DMEM (Agilent cat. no. 103575-100) supplemented with 10mM glucose (Agilent cat. no. 103577-100), 1mM pyruvate (Agilent cat. no. 103578-100) and 2mM glutamine (Agilent cat. no. 103579-100). For mouse vs. human comparisons, MSGN1-Venus+ cells were pre-sorted. Cells were allowed to attach at room temperature for 20 minutes and then transferred to a 37°C incubator without CO_2_ for 1 hour. The Seahorse cartridge was hydrated and calibrated as per the manufacturer instructions. For the Mitochondrial Stress Test (Agilent cat. no. 103015-100), we used oligomycin at 1μM, FCCP at 1μM, rotenone at 0.5μM and antimycin A at 0.5μM. No glucose controls were used to calculate the CO_2_ contribution factor for glycolytic proton efflux rate determination. All samples were run in six to ten replicates in either a Seahorse XF96 or XFe24 Analyzer and the data were analyzed in Wave and Microsoft Excel using macros provided by the manufacturer.

### Extracellular Glucose, Lactate and Glutamine Quantification

Mouse and human MSGN1-Venus+ cells were pre-sorted and seeded onto a fibronectin-coated 96 well plate at a density of 4×10^5^ cells/cm^2^. The media consisted of DMEM (Gibco cat. no. A1443001) containing 2mM glucose, 1mM glutamine, 1mM pyruvate, 0.1mM non-essential amino acids, 1% ITS, 5% dialyzed FBS supplemented (Cytiva cat. no. SH30079.01) with 6 μM Chir 99021, 0.5 μM LDN193189, 50 ng/ml mFgf4, 1 μg/ml Heparin, 2.5 μM BMS493 and 10 μM Rocki. Control samples with media only (no cells) were also included. For glucose and lactate detection, 5μl of media were collected every hour for a total of 6 hours, diluted in 195μl PBS and frozen at −20°C. The Promega Glucose-Glo (Promega cat. no. J6021) and Lactate-Glo (Promega cat. no. J5021) kits were used according to manufacturer protocols on white 384 well plates. Standard curves of glucose and lactate were used to calculate metabolite concentration in the media. For glutamine detection, media was collected at a single timepoint (12 hours) and the Promega Glutamate/Glutamine-Glo kit (Promega cat. no. J8021) was used. Luminescence was measured after the incubation time indicated by the manufacturer using a GloMax Promega plate reader with 1 second integration. Measurements were normalized to cell mass.

### Stable Isotope Tracing

#### Sample preparation

Mouse and human PSM cells were differentiated as described above in 6 well plates. On day 2 of differentiation, the plates were washed once with PBS and replaced with tracer medium. Tracer medium consisted of 25mM either unlabeled or U-^13^C_6_-glucose (Cambridge Isotope Laboratories cat. no. CLM-1396-0.5), 4mM either unlabeled or U-^13^C_5_-glutamine (Cambridge Isotope Laboratories cat. no. CLM-1822-H-0.1), 1mM Sodium Pyruvate, 0.1mM Non-Essential Amino Acids, 100u/ml Penicillin, 100ug/ml Streptomycin, 1% ITS, 5% dialyzed FBS supplemented with 6 μM Chir 99021, 0.5 μM LDN193189, 50 ng/ml mFgf4, 1 μg/ml Heparin, 2.5 μM BMS493 and 10 μM Rocki. Cultures were incubated for 24 hours prior to metabolite extraction. Experiments were performed three times independently.

#### Metabolite extraction

Intracellular metabolites were obtained after washing cells with 2 volumes of room temperature HPLC-grade water and floating the dry plates on liquid nitrogen to quench metabolites. Plates were stored at −80 °C until extraction. Metabolites were extracted with 1 mL 80% MeOH pre-cooled to −80 °C. Insoluble material was removed by centrifugation at 21,000 ×g for 15 min at 4 °C. The supernatant was evaporated to dryness at 42 °C using a SpeedVac concentrator (Thermo Savant). Samples were resuspended in 35 μL LC-MS-grade water prior to analysis.

#### Acquisition parameters

LC-MS analysis was performed on a Vanquish ultra-high-performance liquid chromatography system coupled to a Q Exactive orbitrap mass spectrometer by a HESI-II electrospray ionization probe (Thermo). External mass calibration was performed weekly. Metabolite samples (2.5 μL) were separated using a ZIC-pHILIC stationary phase (2.1 × 150 mm, 5 μm) (Merck). The autosampler temperature was 4 °C and the column compartment was maintained at 25 °C. Mobile phase A was 20 mM ammonium carbonate and 0.1% ammonium hydroxide. Mobile phase B was acetonitrile. The flow rate was 0.1 mL/min. Solvent was introduced to the mass spectrometer via electrospray ionization with the following source parameters: sheath gas 40, auxiliary gas 15, sweep gas 1, spray voltage +3.0 kV for positive mode and −3.1 kV for negative mode, capillary temperature 275 °C, S-lens RF level 40, and probe temperature 350 °C. Data were acquired and peaks integrated using TraceFinder 4.1 (Thermo).

#### Stable isotope quantification

All metabolites except fructose 2,6-bisphosphate (FBP) and 3-phosphoglycerate (3PG) were measured using the following mobile phase gradient: 0 min, 80% B; 5 min, 80% B; 30 min, 20% B; 31 min, 80% B; 42 min, 80% B. The mass spectrometer was operated in selected ion monitoring mode with an m/z window width of 9.0 centered 1.003355-times half the number of carbon atoms in the target metabolite. The resolution was set at 70,000 and AGC target was 1 × 105 ions. Peak areas were corrected for quadrupole bias as previously described [12]. Mass isotope distributions for FBP and 3PG were calculated from full scan chromatograms as described below. Raw mass isotopomer distributions were corrected for natural isotope abundance using a custom R package (mzrtools, https://github.com/oldhamlab/mzrtools) employing the method of Fernandez, et al. [13].

### Cell Volume Measurements

Cells were dissociated in TrypLE, washed and resuspended in PBS. Volume was measured on a Moxi Go II Coulter-principle cell sizer and flow cytometer. When mouse vs. human PSM cells were compared, only MSGN1-Venus+ cells were considered. Data was analyzed in FlowJo.

### Mass and Density Measurements – Suspended Microchannel Resonator

Mouse and human PSM cells were dissociated in TrypLE and MSGN1-Venus+ cells were sorted. Cells were then counted and resuspended in DMEM/F12, 1% ITS, 5% FBS with 6 μM Chir 99021, 20ng/ml bFGF and 0.5 μM LDN193189 at a concentration of 3×10^5^ cells/ml. The cells were kept on ice and their total mass and density were measured using the suspended microchannel resonator (SMR) according a previously developed fluid-switching method [14]. The SMR is a vibrating cantilever with a fluidic channel inside. In the absence of cells, the vibration frequency of the cantilever is proportional to the density of the fluid flowing through the cantilever. As a cell is flown through the cantilever, the vibration frequency of the cantilever changes proportionally to the buoyant mass of the cell. Following the measurements of media density and cell’s buoyant mass in normal media, the cell is immersed in culture media that has been made denser by the addition of 35% OptiPrep (Sigma-Aldrich cat. no. D1556-250ML). The cell is then flow back through the cantilever in the high-density media to obtain a second set of buoyant mass and media density measurements for the cell. The total mass and density of a cell is calculated by comparing these two sets of measurements according to the equation *BM* = *V*(*ρ*_*cell*_ − *ρ*_*fluid*_), where *BM* is buoyant mass, *V* is volume, *ρ*_*cell*_ is density of the cell, and *ρ*_*fluid*_ is density of the fluid. After each cell is measured, the cell is flushed out of the SMR and fresh media and cells are flushed in to the SMR. All measurements were carried out at +4°C. Calibration of the SMR frequency response to a cell was done using NIST traceable 10.12 μm diameter polystyrene beads (Thermo Scientific, Duke Standard beads, cat. no. 4210A), and the calibration of SMR baseline frequency to fluid density was done using NaCl solutions of known density [15].

### Small Molecule Inhibitor Treatments

Human PSM cells were differentiated using the serum-free protocol [2] and treated chronically with the relevant inhibitors or supplements as indicated on Extended Data Table 1 starting on day 2 of differentiation. Timelapse imaging started approximately 2 hours after the inhibitor addition. In the case of aphidicolin, cells were pre-treated for 24 hours before imaging. All other assays (Seahorse, proteasome activity, NAD+/NADH, Peredox fluorescence, etc.) were performed after 16-24 hours of treatment to observe chronic effects.

### Whole-Cell NAD(P)+/NAD(P)H Ratio

Cells were dissociated with TrypLE, washed and resuspended in PBS at a density of 8×10^5^ cells/ml. For each sample, 50μl were distributed in triplicate wells of a 96 well plate. In the case of mouse vs. human comparisons, MSGN1-Venus+ cells were pre-sorted. Samples were lysed by adding 50μl 0.2N NaOH with 1% dodecyltrimethylammonium bromide (DTAB) to preserve the stability of dinucleotides and incubated for 10 minutes at room temperature with gentle shaking. Half of each sample was then transferred to an empty well within the same plate and 25μl 0.4N HCl were added. Samples were then incubated for 15 minutes at 65°C to selectively denature NAD(P)+ in the basic solution and NAD(P)H in the acidic solution. Samples were then cooled down to room temperature for 5 minutes. pH was restored to neutral conditions by adding 25μl 0.5M Trizma® base to acid-treated samples and 50μl Trizma/HCl solution to base-treated samples. 40μl of each sample were then transferred to a white 96 well plate. The detection reagent mixture was prepared according to the instructions in the Promega NAD/NADH (cat. no. G9071) or NADP/NADPH (cat. no. G9081) kit and 40μl were added per sample. Luminescence was measured after an incubation of 45 minutes at room temperature using a GloMax Promega plate reader with 1 second integration. The NAD(P)+/NAD(P)H ratio was calculated as the luminescence ratio of the acid-treated sample to the base-treated sample.

### ADP/ATP Ratio and ATP content measurements

The BioVision ApoSENSOR™ ADP/ATP Ratio Bioluminescent Assay Kit (cat. no. K255-200) was used according to manufacturer’s instructions. Cells were dissociated with TrypLE, washed and resuspended in Nucleotide Releasing Buffer at a density of 3×10^5^ cells/ml. In the case of mouse vs. human comparisons, MSGN1-Venus+ cells were pre-sorted. Background luminescence was measured first, followed by ATP-linked luminescence. To measure ADP, samples were treated with ADP-converting enzyme to generate ATP from ADP. Total (ATP+ADP) luminescence was then recorded. ADP-linked luminescence was calculated by subtracting the ATP-linked luminescence from the total luminescence. A GloMax Promega plate reader with 1 second integration was used.

### Puromycin Incorporation and Pulse-Chase Experiments

To pulse cells with puromycin, samples were washed in PBS and incubated for 1 hour at 37°C 5% CO_2_ in DMEM containing 1μg/ml puromycin (Puro) or 20μM O-propargyl-puromycin (OPP-Puro). Samples were collected immediately following the incubation period for translation rate measurements. For degradation rate measurements, samples were washed with DMEM and chased for the indicated period of time. Specifically, samples were collected every 3 hours for a total of 12 hours. After collection, samples were washed again in PBS and dissociated with TrypLE. One million cells per sample were fixed with 4% formaldehyde and then permeabilized with 0.3% Triton in PBS. Puro-treated samples were then incubated for 1 hour with anti-puromycin Alexa Fluor 647 directly conjugated antibody (1:100; Millipore Sigma cat. no. MABE343). OPP-Puro samples were processed for Click-chemistry detection with Alexa Fluor 647 picolyl azide as per the instructions in the Click-iT® Plus OPP Protein Synthesis Assay Kit (Invitrogen cat. no. C10458). Samples were then analyzed by flow cytometry. In the case of mouse vs. human comparisons, only MSGN1-Venus+ cells were considered.

### L-Azidohomoalanine (AHA) Pulse-Chase Experiments

The Click-it AHA Alexa Fluor 488 Protein Synthesis HCS Assay (Invitrogen cat. no. C10289) was used. Cells were washed in PBS and pre-incubated in DMEM lacking methionine (Gibco cat. no. 21013024 supplemented with 2mM glutamine, 1mM sodium pyruvate, 1% non-essential amino acids and 0.2mM cysteine) for one hour prior to the AHA pulse. The medium was then replaced by methionine-free DMEM containing 50μM AHA and cells were incubated for one hour. Samples were either processed immediately or after the indicated chase time as per the manufacturer instructions. Samples were then analyzed by flow cytometry

### Proteasome Activity Assays

The Promega Proteasome-Glo Chemotrypsin-like assay (Promega cat. no. G8660) was used following the manufacturer’s protocols. Cells were dissociated with TrypLE, washed several times and resuspended in DMEM at a density of 8×10^5^ cells/ml. In the case of mouse vs. human comparisons, MSGN1-Venus+ cells were pre-sorted. Luminescence was recorded on a GloMax Promega plate reader with 1 second integration.

### Statistical Analyses

Statistical analyses were performed with Prism 9 software (GraphPad). P values <0.05 were considered significant. Details of statistical analyses are indicated in figure legends. Unpaired t-tests or ordinary one-way ANOVA were performed with Tukey correction for multiple comparisons. All differentiation experiments were performed a minimum of three independent times (rounds of differentiation), each containing at least three technical replicates (wells) per condition.

